# *Leishmania* secretory factor GP63 targets the DICER1/miR-122/hepcidin axis in host macrophages to deplete Nramp1

**DOI:** 10.1101/2025.04.16.649108

**Authors:** Suman Samanta, Sourav Banerjee, Rupak Datta

## Abstract

Micronutrient sequestration is a powerful host defense against intracellular pathogens. A key player in this is Nramp1, which effluxes iron from phagolysosomes, depriving the engulfed pathogens of this essential element. *Leishmania* counters this by triggering hepcidin-mediated proteasomal degradation of Nramp1. Interestingly, *Leishmania major* conditioned media induced hepcidin expression and Nramp1 degradation even in uninfected macrophages, resulting in enhanced endo/lysosomal iron. This observation led to the identification of the secretory factor GP63 responsible for Nramp1 degradation. Conditioned media from the LmGP63^−/-^ strain failed to upregulate hepcidin or degrade Nramp1. Further, GP63 was found to deplete macrophage DICER1, impairing maturation of miR-122, a negative regulator of hepcidin. Consistent with *in vitro* results, the LmGP63^−/-^ strain, unlike its wild type counterpart, was unable to deplete DICER1, induce hepcidin expression or suppress Nramp1 in infected mice. Collectively, we uncover a novel role for *Leishmania* GP63 in targeting the host DICER1/miR-122 axis to trigger hepcidin expression and Nramp1 degradation, facilitating iron acquisition by the parasite.

**SUMMARY:** This study uncovers a novel role for the *Leishmania* secretory protein GP63 in targeting the DICER1/miR-122 axis in host macrophages to upregulate hepcidin and promote Nramp1 degradation. By employing this strategy, *Leishmania* parasites boost iron availability in their replication niche.

## INTRODUCTION

*Leishmania* spp. are protozoan parasites of medical significance belonging to the Trypanosomatidae family. They cause a spectrum of diseases collectively known as leishmaniasis, with symptoms ranging from self-healing cutaneous lesions to potentially fatal visceral infection, known as kala-azar (Burza et al., 2018). With an estimated 0.7 to 1 million new cases occurring annually, leishmaniasis continues to be a significant global health burden (Burza et al., 2018; Alvar et al., 2012). Lack of a vaccine and increasing resistance to the few existing drugs highlight the need for better understanding of the disease pathogenesis, which could lead to the development of alternative therapies (Ponte-Sucre et al., 2017; Hefnawy et al., 2017).

During its dimorphic lifecycle, *Leishmania* alternates between a sand fly vector and its mammalian host. In the midgut of the sand fly, the parasite proliferates in its promastigote form before being transmitted to the host during a blood meal. Following transmission, *Leishmania* promastigotes are rapidly engulfed by host macrophages where they initially remain entrapped within a phagosome, which gradually matures into acidic phagolysosome. Exposure to the phagolysosomal environment drives transformation of long, flagellated, and highly motile promastigotes into rounded amastigotes with short, non-motile flagella (Chang and Dwyer, 1978; Peters et al., 2008). Apart from being acidic, this free radical-rich phagolysosomal compartment has limited nutrient availability, posing a significant challenge to the survival of intracellular *Leishmania* (Haas, 2007; McConville et al., 2007). How the parasite hijacks host cell machinery to metabolically adapt to such adverse conditions remains to be fully understood. In this context, investigating *Leishmania’*s iron acquisition strategy within the phagolysosomal niche is of utmost importance.

Iron is an essential micronutrient for all life forms, including *Leishmania*. It acts as a cofactor in several key metalloenzymes and is therefore crucial for the proliferation of intracellular parasites (Cairo et al., 2006; Taylor and Kelly, 2010). Thus, it is not surprising that *Leishmania amazonensis* lacking the ferrous iron transporter LIT1 failed to replicate within macrophages and lost its ability to cause infection in mice (Huynh et al., 2006). However, despite its indispensability, free iron is highly toxic due to its participation in the Fenton reaction, which generates reactive hydroxyl radicals (Dixon and Stockwell, 2014). Therefore, iron availability is tightly regulated in mammalian cells and intracellular *Leishmania* must overcome macrophage’s nutritional immunity to access host iron pool (Nairz et al., 2010; Flannery et al., 2013). Macrophage iron content is controlled by the interplay of ferritin and ferroportin. Ferritin stores excess iron in a non-toxic, bioavailable form in the cytosol, while ferroportin, located on the plasma membrane exports iron out of the cell to maintain cellular iron homeostasis (Harrison and Arosio, 1996; Drakesmith et al., 2015). Iron uptake in macrophages occurs through two main pathways: transferrin-receptor-mediated endocytosis and phagocytosis of senescent RBCs, both delivering iron to the lysosome/phagolysosome for recycling (Muckenthaler et al., 2017; Alford et al., 1991). Of particular interest is the natural resistance-associated macrophage protein 1 (Nramp1, also known as SLC11A1), an iron exporter located on the endo/lysosomal membrane that effluxes iron from the phagolysosomes, limiting its availability in this compartment (Vidal et al., 1993; Gruenheid et al., 1997; Barton et al., 1999).

The *Nramp1* gene was originally identified through positional cloning as a key determinant of resistance against unrelated intracellular pathogens like *Mycobacteria*, *Salmonella* and *Leishmania* (Vidal et al., 1993, 1995). A naturally occurring G169D point mutation in the transmembrane domain of Nramp1 disrupts its proper maturation, rendering inbred mouse strains carrying this mutation more susceptible to these infections (Vidal et al., 1993, 1996). While Nramp1 is primarily expressed in phagocytic cells of the myeloid lineage, particularly macrophages, it has also been detected in neurons and microglial cells of the mouse brain (Govoni et al., 1997; Evans et al., 2001). Nramp1 co-localizes with late endosomal/lysosomal markers in macrophages and is also recruited to the maturing phagosome (Gruenheid et al., 1997; Searle et al., 1998; Cheng and Wang, 2012). Studies in *Dictyostelium* and mammalian macrophage cell have established the role of Nramp1 in mediating efflux of ferrous iron from the phagosome (Barton et al., 1999; Atkinson and Barton, 1999; Buracco et al., 2015). This facilitates efficient recycling of haemoglobin-derived iron in macrophages (Soe-Lin et al., 2008). In addition to iron, Nramp1 has also been shown to extrude manganese from the phagosomal compartment in a pH-dependent manner (Jabado et al., 2000). Another study demonstrated that Nramp1 restricts *Salmonella* growth by depriving it of magnesium (Cunrath and Bumann, 2019). Thus, it has been proposed that Nramp1 confers nutritional immunity by limiting the availability of iron and other divalent metals to invading pathogens (Marquis and Gros, 2007).

Despite its critical role in determining the outcome of host-pathogen interaction, whether Nramp1 levels are modulated during infection remained unknown until recently. We investigated this in *Leishmania major* infected macrophages and observed that at 12 hours post-infection, Nramp1 levels were significantly reduced but were restored to normalcy by 30 hours (Banerjee and Datta, 2020). Consistent with its role as a phagosomal iron exporter, Nramp1 depletion was associated with increased phagolysosomal iron content and a higher intracellular parasite burden. Further, we showed that this infection-induced depletion of Nramp1 was caused by ubiquitin-proteasomal degradation, mediated by its interaction with the iron-regulatory peptide hormone hepcidin, whose expression was markedly upregulated during infection (Banerjee and Datta, 2020). It was evident from these results that *Leishmania* parasite targets Nramp1 to inhibit iron efflux from its replication niche, the phagolysosome. However, the key question remained: how does *Leishmania* infection trigger hepcidin expression and target Nramp1 for proteasomal degradation? Taking cue from an intriguing observation that Nramp1 depletion was observed even in bystander uninfected cells in close proximity to *Leishmania* infected macrophages, we hypothesized that a parasite-secreted factor might be driving this process. To test this, we treated macrophages with *Leishmania* conditioned media, which induced proteasomal degradation of Nramp1 to a similar extent as direct infection. This discovery set off a series of experiments that culminated in the identification of GP63 as the causative secretory factor. Compelling evidence for this came from the observation that conditioned medium from GP63-deficient parasites failed to cause Nramp1 degradation. GP63 is a zinc-dependent metalloprotease of *Leishmania* that facilitates immune evasion by targeting multiple host proteins; however, until now it has not been reported to target any iron transporter in macrophage (Guay-Vincent et al., 2022; Isnard et al., 2012). Our *in vitro* and *in vivo* animal infection experiments collectively reveal that by depleting DICER1/miR-122, GP63 drives hepcidin upregulation in macrophages, leading to proteasomal degradation of Nramp1. This unique strategy enables *Leishmania* to subvert host iron homeostasis for its own benefit.

## RESULTS

### Treatment of macrophage cells with *Leishmania* conditioned media results in proteasomal degradation of Nramp1

Recently, we reported that *L. major* infection in macrophages caused a significant reduction in Nramp1 protein levels at 12 hours post-infection. This correlated with a simultaneous increase in phagolysosomal iron content and a higher intracellular parasite burden (Banerjee and Datta, 2020). Since our infection protocol routinely yields 65-70% infection rate, we were curious to analyse the Nramp1 levels in the bystander uninfected macrophages surrounded by the *Leishmania* infected ones. Consistent with our previous findings, *L. major* infection in J774A.1 macrophages resulted in overall ∼ 2 folds reduction in Nramp1 protein levels, as evidenced by the immunofluorescence results (Fig. 1, A and B). Careful images analysis revealed that even the bystander uninfected cells (Lm-) exhibited a similar reduction in Nramp1 levels as their neighbouring infected macrophages (Lm+) (Fig. 1, A inset and B; and Fig. S1A). This intriguing data suggested that a secretory factor of the parasite could be involved in reducing the Nramp1 levels in macrophage cells. To investigate whether a secretory factor of *Leishmania* can indeed cause reduction of Nramp1 levels in host macrophages, we cultured *L. major* promastigotes, collected the conditioned media (Lm-CM) containing the parasite’s secretory factors and treated macrophage cells with it (Fig. 1C). By microscopic observation and western blot analysis using antibodies against known cytosolic proteins of *Leishmania*, we confirmed that the collected Lm-CM was devoid of any intact parasite or their cytosolic content (Fig. S1, B and C). Treatment of J774A.1 macrophages with Lm-CM for 12 hours led to ∼ 2 folds reduction in the Nramp1 levels, as demonstrated by immunofluorescence microscopy (Fig.1, D and E). This intriguing finding was validated in primary peritoneal macrophages derived from BALB/c mice, which also showed a similar reduction in the Nramp1 levels upon Lm-CM treatment (Fig. S1, D and E). qRT-PCR data demonstrated that Lm-CM-mediated reduction in Nramp1 levels was not due to transcriptional downregulation of Nramp1 (Fig. S1F). However, as revealed by our immunofluorescence and western blot data, depletion of Nramp1 could be prevented by treating macrophage cells with the proteasome inhibitor MG132, suggesting involvement of the proteasomal degradation pathway in the process (Fig. 1, F and G; and Fig. S1, G and H). In agreement with our previous finding with *L. major* infection, we observed ∼ 1.7-folds increase in the endo/lysosomal iron content in the Lm-CM-treated J774A.1 macrophages compared to the untreated control (Fig. 1H). Interestingly, from our western blot data it was evident that Lm-CM lost its ability to cause Nramp1 degradation when it was pre-heated at 95^0^C or pre-incubated with trypsin (Fig. 1, I and J). These results suggests that a heat and protease-sensitive secreted factor (most likely a protein) of the parasite is responsible for Nramp1 degradation in host macrophages.

**Figure 1.**
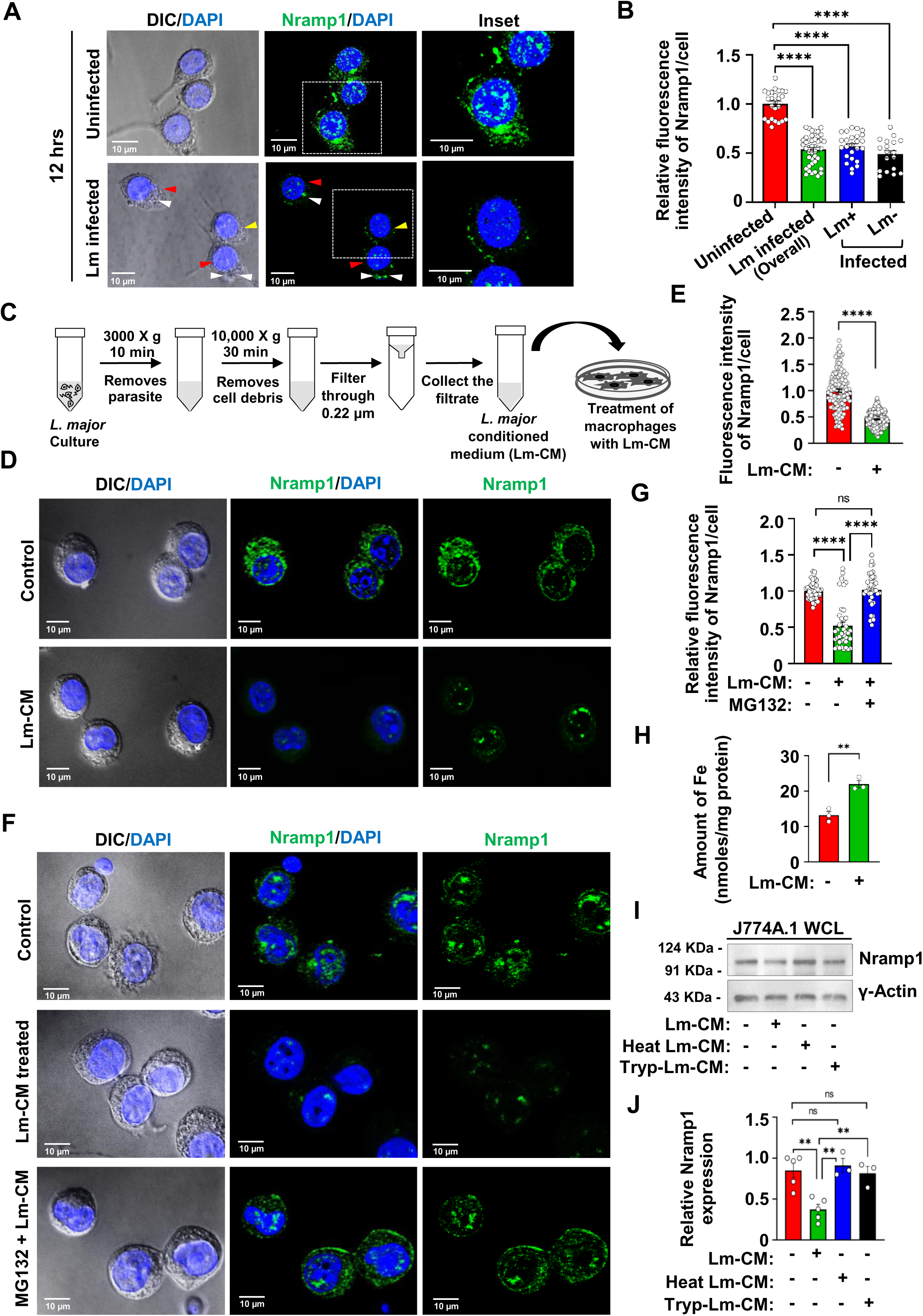
*L. major* conditioned media treatment depleted Nramp1 and elevated endo/lysosomal iron levels in macrophages. **(A)** Nramp1 was visualised by immunostaining with anti-Nramp1 (green) in uninfected or *L. major* (Lm)-infected J774A.1 macrophages. Nuclei were stained with DAPI (blue). DIC/DAPI panel shows the presence of intracellular parasites (smaller nuclei, indicated by white arrows) in infected cells (marked by red arrows). Yellow arrows mark the uninfected cells. The insets are zoomed regions marked by boxes. Images were acquired with Leica SP8 confocal, 63× objective. **(B)** Quantification of the Nramp1 fluorescence intensities from uninfected and Lm-infected (overall) macrophages is shown in bar diagram. For the Lm-infected macrophages, fluorescence intensities were separately analyzed for bystander uninfected cells (Lm-) and the infected cells (Lm+). The data are expressed as means ± SEM (Uninfected, n= 25; *L. major* infected (overall), n= 30; Lm+, n= 25; Lm-, n= 18 cells from N = 3 independent experiments). **(C)** Schematic illustration showing the preparation of *L. major* conditioned medium (Lm-CM). **(D)** Nramp1 was visualised by immunostaining with anti-Nramp1 (green) in J774A.1 macrophages treated for 12 hours with *L. major* conditioned medium (Lm-CM) or with M199 medium only (control). Nuclei were stained with DAPI (blue) and the DIC/DAPI panel shows the overall cell morphology. Images were acquired with Carl Zeiss Apotome.2 microscope, 63× objective. **(E)** Quantification of the Nramp1 fluorescence intensities in the respective images shown in bar diagram. The data are expressed as means ± SEM (at least 150 cells from N = 3 independent experiments were analyzed). **(F)** Nramp1 immunostaining (green) in J774A.1 macrophages treated for 12 hours with just M199 media (control) or with Lm-CM or Lm-CM + 1µm MG132 (macrophages were pre-treated with MG132 prior to Lm-CM treatment). Nuclei were stained with DAPI (blue) and the DIC/DAPI panel shows the overall cell morphology. Images were acquired with Carl Zeiss Apotome.2 microscope, 63× objective. **(G)** Quantification of the Nramp1 fluorescence intensities in the respective images shown in bar diagram. Values expressed as means ± SEM (at least 40 cells from N = 3 independent experiments were analyzed). **(H)** Quantification of endo/lysosomal iron in J774A.1 macrophages treated for 12 hours with just M199 medium (-) or with Lm-CM (+). The data are presented as means ± SEMs from three independent experiments. **(I)** Representative western blots of Nramp1 and Ɣ-actin (loading control) on J774A.1 macrophage whole cell lysates (WCL) prepared from either control cells (treated just with M199 media) or cells treated for 12 hours with Lm-CM, heat-inactivated Lm-CM or trypsinized Lm-CM. **(J)** Bar diagram showing the quantification of Nramp1 band densities in the respective samples normalized to Ɣ-actin. Values expressed as means ± SEMs from at least three independent experiments. In all bar diagrams, individual values are shown as small circles. n.s., non-significant; ****P ≤ 0.0001, **P ≤ 0.01, estimated by two-tailed unpaired Student’s t-test.

### Treatment of *Leishmania* conditioned media with metalloprotease inhibitors abrogated its ability to cause Nramp1 degradation

*Leishmania* secretes a wide variety of proteins either in free form, or packed within exosomal vesicles (EVs) (Santarém et al., 2013; Pissarra et al., 2022). So, we were curious to know whether this unknown secretory protein of *L. major* responsible for Nramp1 degradation is released in its free form or bound to exosomes. Using an exosome isolation kit, we isolated the EVs from the Lm-CM and suspended them in M199 media. Alongside, the post-EV supernatant was also collected (Fig. 2A). Authenticity of both the fractions was confirmed by scanning electron microscopy (SEM) (Fig. 2B). Interestingly, our western blot and immunofluorescence data demonstrated that treatment of J774A.1 macrophages with either EVs or the post-EV supernatant fraction resulted in a significant reduction in Nramp1 levels, similar to the effects observed with Lm-CM treatment (Fig. 2, C - F). Exoproteome analysis of *L. infantum* revealed that only a limited number of secreted proteins are present in both exosomal and non-exosomal fractions, with GP63, a zinc-dependent metalloprotease, being the most abundant among them (Santarém et al., 2013). Taking cue from this, we analyzed the potential role of GP63 in Nramp1degradation by pharmacological inhibition approach. For this, we pre-incubated the Lm-CM with the zinc-chelating metalloprotease inhibitors EDTA or 1,10-Phenanthroline and then treated the macrophage cells with it (Chaudhuri et al., 1989). Our immunofluorescence as well as western blot data showed that pre-incubation with EDTA or 1,10-Phenanthroline abrogated Lm-CM’s ability to cause Nramp1 degradation. However, exogenous addition of zinc restored this ability in EDTA/1,10-Phenanthroline-treated Lm-CM (Fig. 3, A - F). It may be noted that treatment of macrophage cells with EDTA, 1,10-Phenanthroline or ZnCl_2_ alone did not cause any change in the Nramp1 levels (Fig. S2, A and B). These results suggest that the zinc-chelator-sensitive GP63 present in Lm-CM may be responsible for Nramp1 degradation in macrophages.

**Figure 2.**
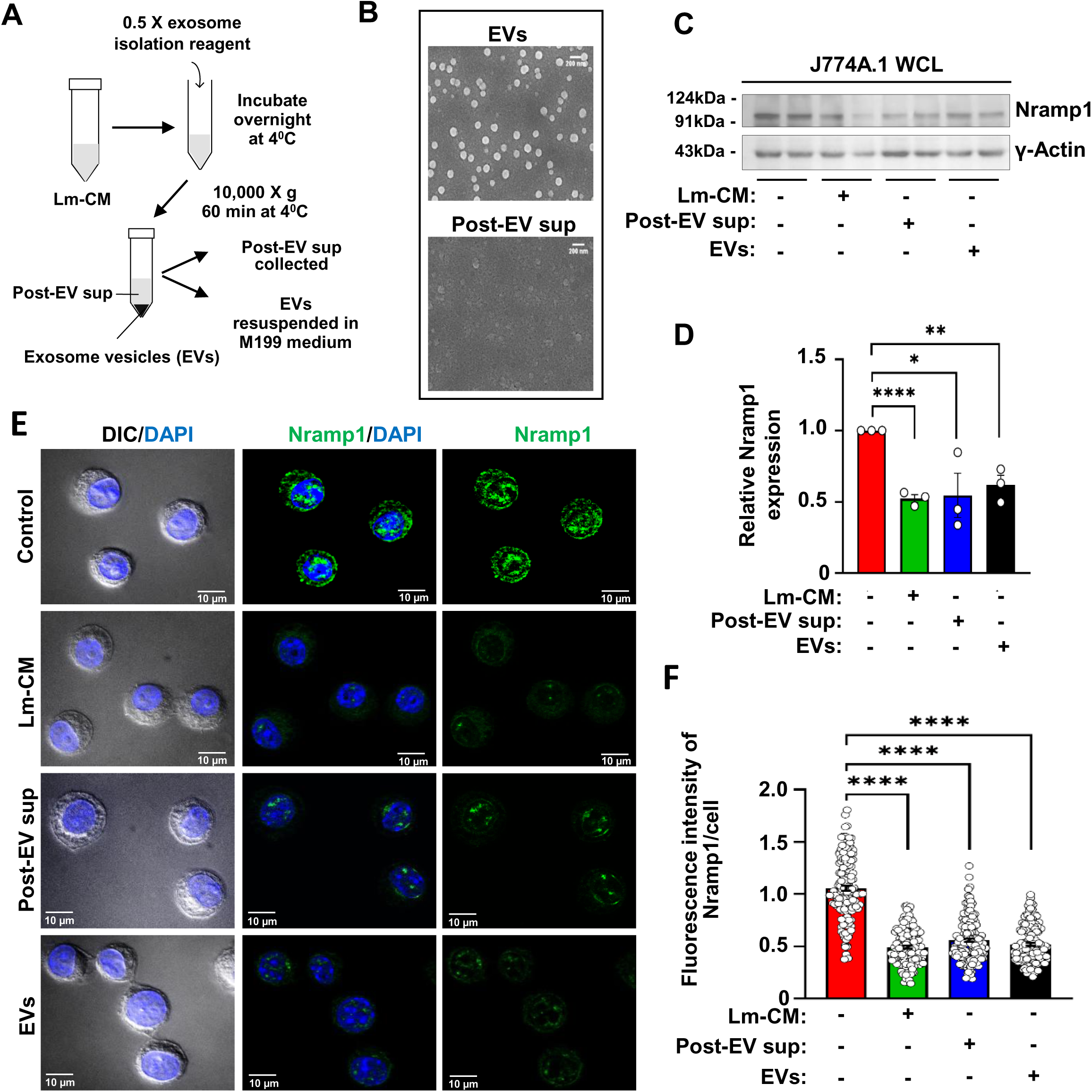
Treatment with exosomal and non-exosomal fractions of *L. major* conditioned media depleted Nramp1 levels in macrophages. **(A)** Schematic illustration of isolation of exosomal vesicles (EVs) from Lm-CM. **(B)** Representative SEM images of EV fraction (upper image) and post-EV supernatant fraction (lower image) isolated from Lm-CM. Scale bars, 200 nm. **(C)** Representative western blots of Nramp1 and Ɣ-actin (loading control) on J774A.1 macrophage whole cell lysates (WCL) obtained from control macrophages (treated just with M199 media) or macrophages incubated for 12 hours with Lm-CM, post-EV supernatant or EVs fraction. **(D)** Bar diagram showing the quantification of Nramp1 band densities in the respective samples normalized to Ɣ-actin. Values expressed as means ± SEMs from three independent experiments. Values expressed as means ± SEMs from at least three independent experiments. **(E)** Nramp1 immunostaining (green) in J774A.1 macrophages treated for 12 hours with just M199 media (control) or Lm-CM, post-EV supernatant or EVs. Nuclei were stained with DAPI (blue) and the DIC/DAPI panel shows the overall cell morphology. Images were acquired with Carl Zeiss Apotome.2 microscope, 63× objective. **(F)** Quantification of the Nramp1 fluorescence intensities in the respective images shown in bar diagram. Values are expressed as means ± SEMs (at least 147 cells from N = 3 independent experiments were analyzed). In all bar diagrams, individual values are shown as small circles. ****P ≤ 0.0001, **P ≤ 0.01, *P ≤ 0.05, estimated by two-tailed unpaired Student’s t-test.

**Figure 3.**
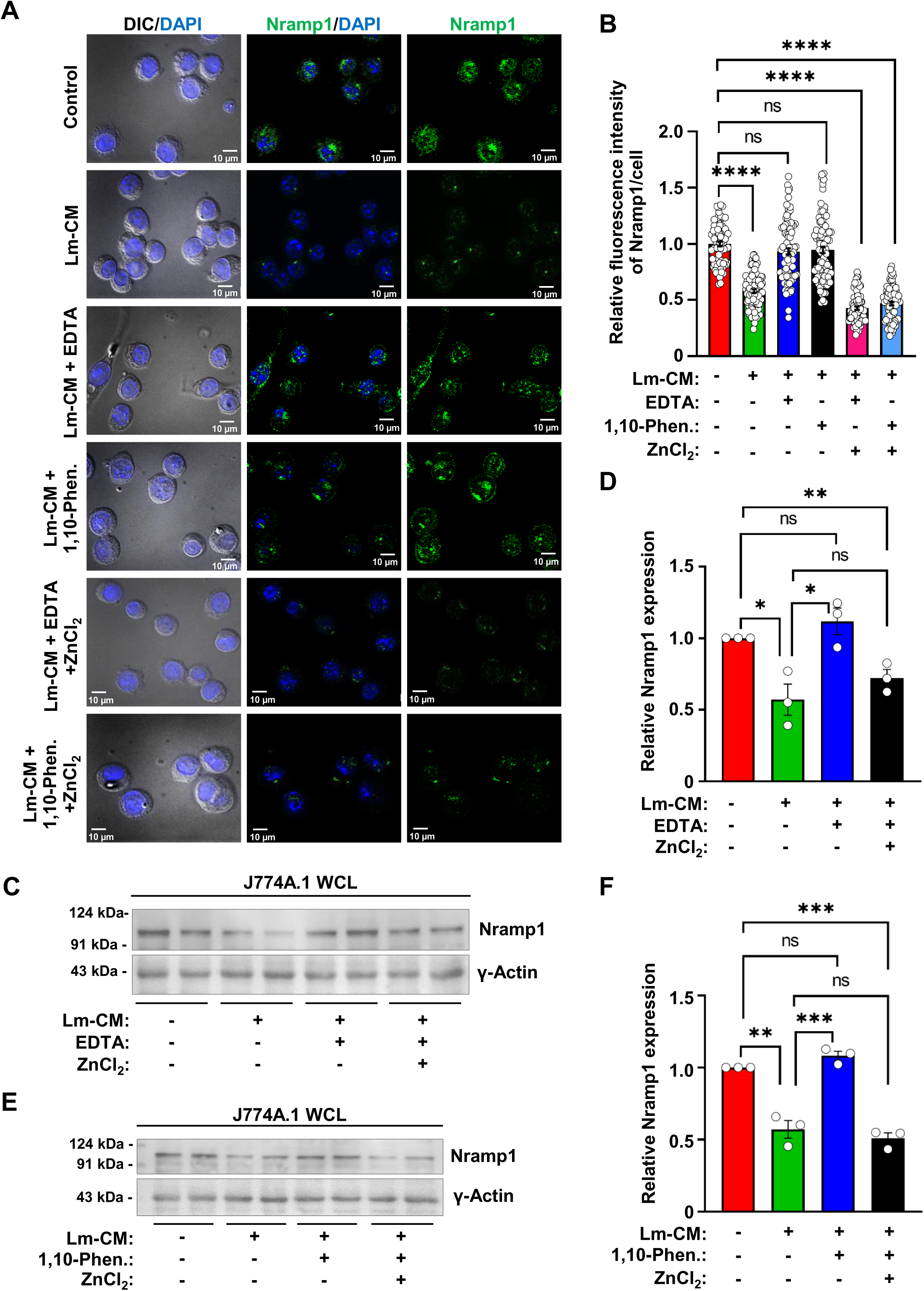
Treatment with metalloprotease inhibitors abrogated the ability of *L. major* conditioned media to cause Nramp1 depletion. **(A)** Nramp1 immunostaining (green) in J774A.1 macrophages treated for 12 hours with just M199 media (control) or Lm-CM, Lm-CM + 1 mM EDTA, Lm-CM + 1 mM 1,10-Phenanthroline, Lm-CM + 1 mM EDTA + 1mM ZnCl_2_, Lm-CM + 1 mM 1,10-Phenanthroline + 1mM ZnCl_2_. Nuclei were stained with DAPI (blue) and the DIC/DAPI panel shows the overall cell morphology. Images were acquired with Carl Zeiss Apotome.2 microscope, 63× objective. **(B)** Quantification of the Nramp1 fluorescence intensities in the respective images shown in bar diagram. Values are expressed as means ± SEMs (at least 75 cells from N = 3 independent experiments were analyzed). **(C)** Representative western blots of Nramp1 and Ɣ-actin (loading control) on J774A.1 macrophage whole cell lysates (WCL) obtained from untreated macrophages (treated just with M199 media) or macrophages treated for 12 hours with Lm-CM or Lm-CM + 1mM EDTA or Lm-CM + 1mM EDTA + 1mM ZnCl_2_. **(D)** Bar diagram showing the quantification of Nramp1 band densities in the respective samples normalized to Ɣ-actin. Values expressed as means ± SEMs from at least three independent experiments. **(E)** Representative western blots of Nramp1 and Ɣ-actin (loading control) on J774A.1 macrophage whole cell lysates (WCL) obtained from untreated macrophages (treated just with M199 media) or macrophages treated for 12 hours with Lm-CM or Lm-CM + 1mM 1,10-Phenanthroline or Lm-CM + 1mM 1,10-Phenanthroline + 1mM ZnCl_2_. **(F)** Bar diagram showing the quantification of Nramp1 band densities in the respective samples normalized to Ɣ-actin. Values expressed as means ± SEMs from at least three independent experiments. In all bar diagrams, individual values are shown as small circles. n.s., non-significant; ****P ≤ 0.0001,***P ≤ 0.001,**P ≤ 0.01, *P ≤ 0.05 estimated by two-tailed unpaired Student’s t-test.

### Conditioned media from the GP63^−/-^ *Leishmania* failed to cause Nramp1 degradation

To obtain definitive evidence that GP63 is indeed responsible for degradation of Nramp1, we planned to generate a GP63 knockout *L. major* strain (LmGP63^−/-^) using CRISPR-Cas9. According to the *L. major* genome data base, four copies of the GP63 gene are arranged in tandem on Chromosome 10 (Ivens et al., 2005). On the background of an engineered *L. major* strain stably expressing Cas9 and T7 RNA polymerase (LmCas9/T7), we aimed to knock out both the alleles of all four copies of GP63 using sgRNAs that specifically targeted the 5’ UTR of GP63-1 and 3’ UTR of GP63-4 (Fig. S3A, Fig. 4, A and B). Puromycin and blasticidin repair cassettes replaced the cleaved target sequences, resulting in generation of the LmGP63^−/-^ strain, which was selected in the presence of puromycin dihydrochloride and blasticidin S hydrochloride. The authenticity of the LmGP63^−/-^ strain was verified by genomic PCR using appropriate primers sets confirming the presence of puromycin (600 bp product with primers P1/P2) and blasticidin (390 bp product with primers P3/P4) cassettes and absence of the GP63 gene (checked with primers P5/P6) (Fig. S3, B and C). Complete absence of the GP63 protein in the LmGP63^−/-^ strain was confirmed by immunofluorescence (Fig. 4C), flowcytometry (Fig. S3D) and gelatin zymography assay (Fig. 4D). Next, we collected the conditioned media from the LmGP63^−/-^ strain (LmGP63^−/-^-CM) and also from two other control strains, wild type *L. major* (Lm-CM) and LmCas9/T7 (LmCas9/T7-CM). Macrophage cells were treated with these three conditioned media in parallel to compare their abilities to cause Nramp1 degradation. Our immunofluorescence (Fig. 4, E and F) and western blot (Fig. 4, G and H) data showed that while Lm-CM and LmCas9/T7-CM could induce Nramp1 degradation in macrophages, LmGP63^−/-^-CM completely lacked this ability. These results provided a clear evidence that GP63 in Lm-CM is responsible for inducing proteasomal degradation of Nramp1 in host macrophages and prompted us to investigate the underlying mechanism.

**Figure 4.**
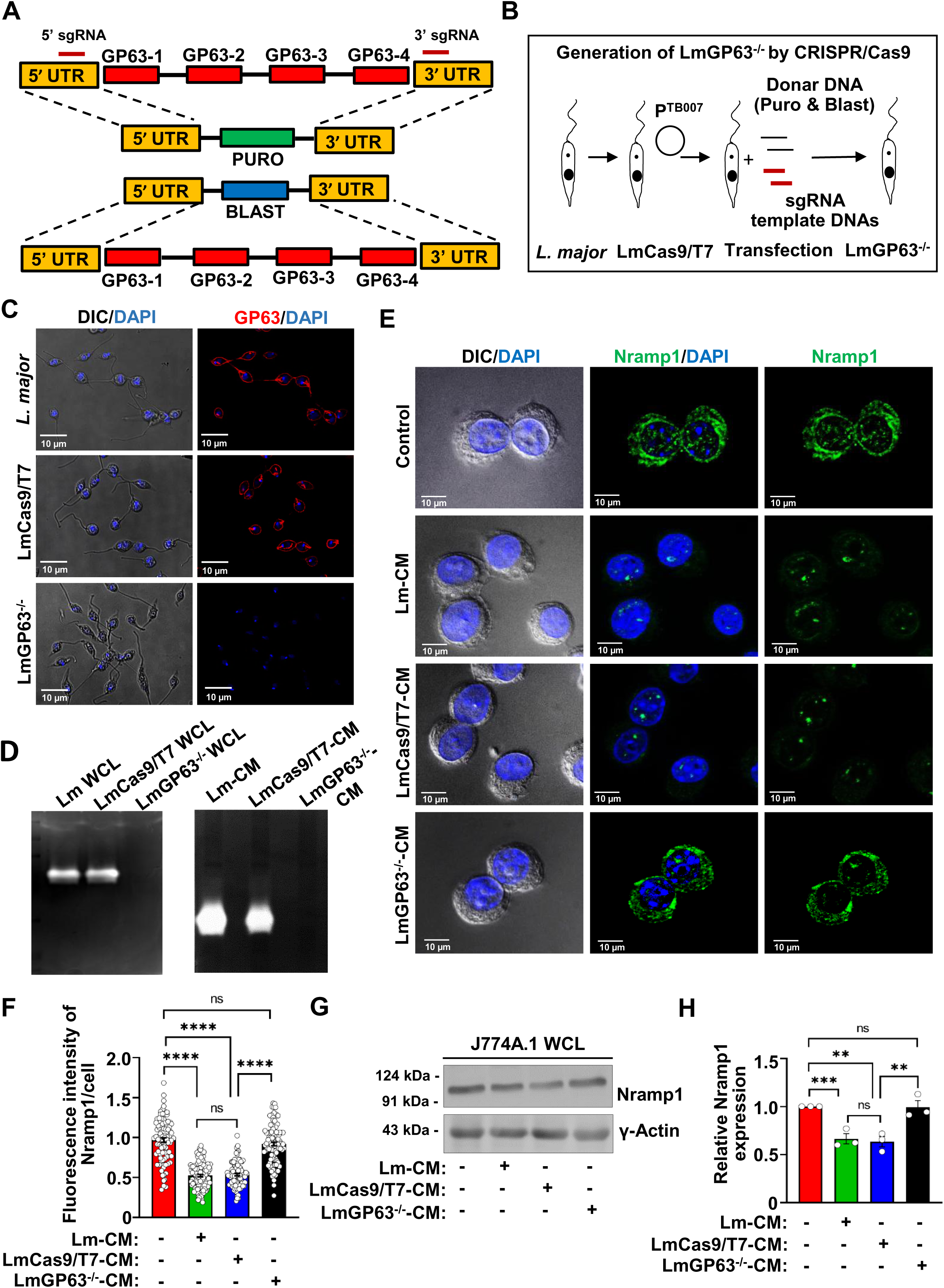
Conditioned media from the GP63 knockout *L. major* (LmGP63^−/-^) failed to deplete Nramp1 in macrophages. **(A and B)** Schematic illustrations of the GP63 locus in the *L. major* genome and the strategy adopted to generate LmGP63^−/-^ strain using CRISPR/Cas9 technique. **(C)** GP63 immunostaining (red) with anti-GP63 in wild type *L. major*, LmCas9/T7, and LmGP63^−/-^ promastigotes. Nuclei were stained with DAPI (blue) and the DIC/DAPI panel shows the overall cell morphology. Images were acquired with Leica SP8 confocal, 63× objective. **(D)** Representative gelatin zymograms showing the absence of GP63 activity (absence of white bands, indicating gelatin proteolysis) in LmGP63^−/-^ WCL compared to *L. major* WCL and LmCas9/T7 WCL (left zymogram) and in conditioned media from the LmGP63^−/-^ strain (LmGP63^−/-^-CM) compared to Lm-CM and LmCas9/T7-CM (right zymogram). **(E)** Nramp1 was visualised by immunostaining with anti-Nramp1 (green) in J774A.1 macrophages treated for 12 hours with M199 media only (control) or with Lm-CM, LmCas9/T7-CM or LmGP63^−/-^-CM. Nuclei were stained with DAPI (blue) and the DIC/DAPI panel shows the overall cell morphology. Images were acquired with Carl Zeiss Apotome.2 microscope, 63× objective. **(F)** Quantification of the Nramp1 fluorescence intensities in the respective images shown in bar diagram. Values are expressed as means ± SEMs (at least 90 cells from N = 3 independent experiments were analyzed). **(G)** Representative western blots of Nramp1 and Ɣ-actin (loading control) on J774A.1 macrophage whole cell lysates (WCL) obtained from untreated macrophages (treated just with M199 media) or macrophages treated for 12 hours with Lm-CM, LmCas9/T7-CM or LmGP63^−/-^-CM. **(H)** Bar diagram showing the quantification of Nramp1 band densities in the respective samples normalized to Ɣ-actin. Values are expressed as means ± SEM from 3 independent experiments. In all bar diagrams, individual values are shown as small circles. n.s., non-significant; ****P ≤ 0.0001, ***P ≤ 0.001,**P ≤ 0.01 estimated by two-tailed unpaired Student’s t-test.

### Conditioned media from the GP63^−/-^ *Leishmania* lost the ability to induce hepcidin expression or trigger ubiquitination of Nramp1 in host macrophages

Previously, we reported that infection of macrophage cells with *L. major* led to transcriptional upregulation of hepcidin, which in-turn promoted ubiquitination and proteasomal degradation of Nramp1 (Banerjee and Datta, 2020). Since conditioned media from the GP63^−/-^ *L. major* was unable to cause proteasomal degradation of Nramp1, we sought to determine whether this is due to its inability to induce hepcidin expression or trigger ubiquitination of Nramp1. Consistent with our previous findings in *L. major*-infected macrophages, we found that macrophage cells treated with Lm-CM or LmCas9/T7-CM exhibited significant upregulation of hepcidin at both protein (Fig. 5, A - C) and mRNA levels (Fig. 5D), along with increased Nramp1 ubiquitination (Fig. 5, E and F). Interestingly, treatment with LmGP63^−/-^-CM neither led to hepcidin upregulation nor caused Nramp1 ubiquitination (Fig. 5, A-F). This suggest that GP63 in Lm-CM plays a pivotal role in promoting hepcidin expression, ultimately driving Nramp1 ubiquitination and its proteasomal degradation. Thus, our next goal was to investigate the mechanism by which GP63 regulates hepcidin expression.

**Figure 5.**
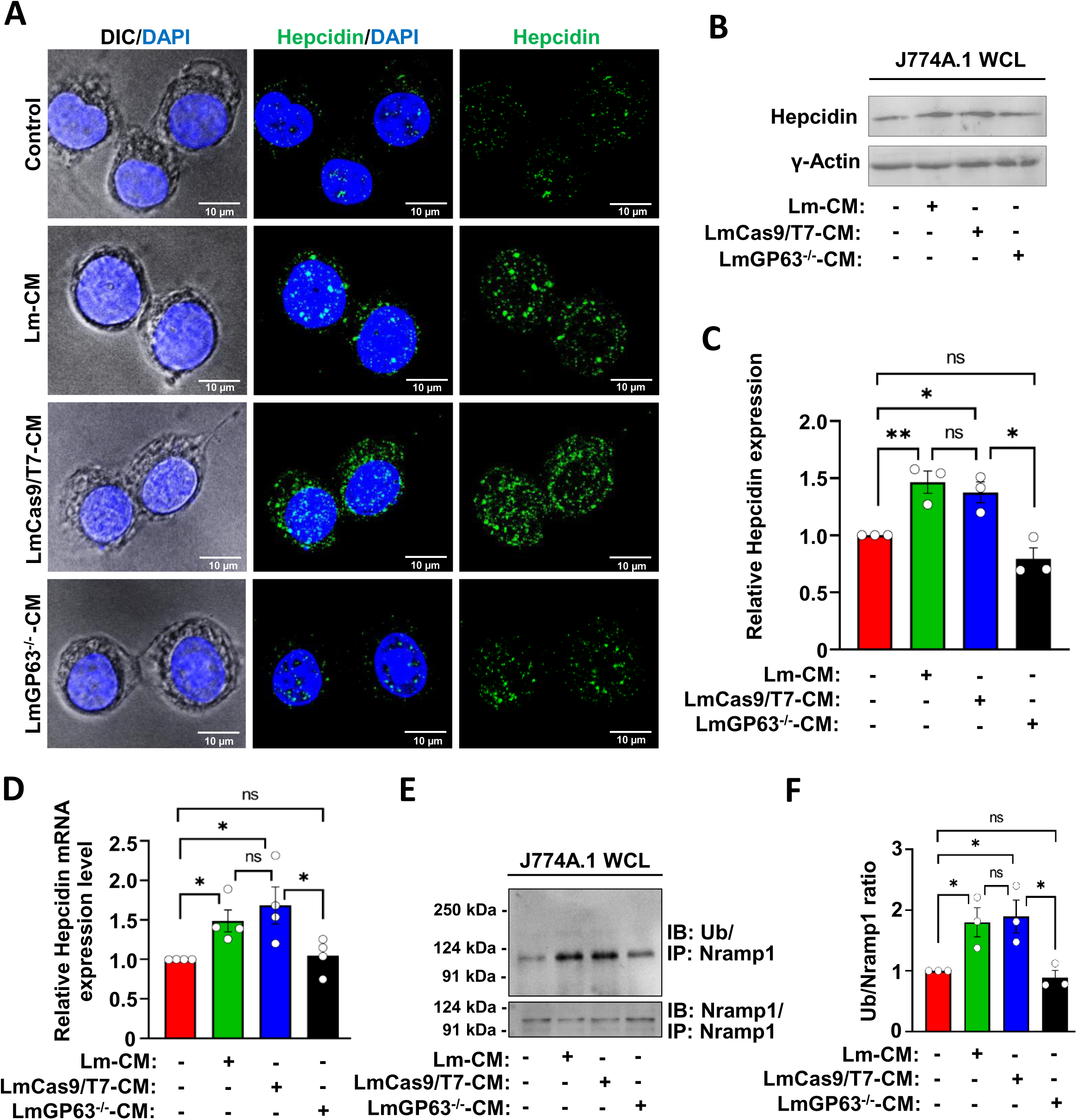
Conditioned media from the LmGP63^−/-^ strain unable to induce hepcidin expression or trigger Nramp1 ubiquitination in macrophages. **(A)** Hepcidin was visualised by immunostaining with anti-hepcidin (green) in J774A.1 macrophages treated for 12 hours with M199 media only (control) or with Lm-CM, LmCas9/T7-CM or LmGP63^−/-^-CM. Nuclei were stained with DAPI (blue) and the DIC/DAPI panel shows the overall cell morphology. Images were acquired with Carl Zeiss Apotome.2 microscope, 63× objective. **(B)** Representative western blots of hepcidin and Ɣ-actin (loading control) on J774A.1 macrophage whole cell lysates (WCL) obtained from untreated macrophages (treated just with M199 media) or macrophages treated for 12 hours with Lm-CM, LmCas9/T7-CM or LmGP63^−/-^-CM. **(C)** Bar diagram showing the quantification of hepcidin band densities in the respective samples normalized to Ɣ-actin. Values are expressed as means ± SEMs from 3 independent experiments. **(D)** Bar diagram showing qRT-PCR data of relative hepcidin expression in J774A.1 macrophages treated for 12 hours with M199 media only (control) or with Lm-CM, LmCas9/T7-CM or LmGP63^−/-^-CM. All the measurements were performed using the control cell as reference sample (expression level set to 1.0) and β-actin as an endogenous control gene for normalization. Values are expressed as means ± SEMs from N = 4 independent experiments. **(E)** Immunoprecipitation (IP) of Nramp1 from J774A.1 macrophage whole cell lysates (WCL) obtained from untreated macrophages (treated just with M199 media) or macrophages treated for 12 hours with Lm-CM, LmCas9/T7-CM or LmGP63^−/-^-CM, followed by probing with anti-ubiquitin antibody (top panel) or with anti-Nramp1 antibody (bottom panel). **(F)** Bar diagram showing the quantification of ubiquitin band densities in the respective samples normalized to Nramp1. Values are expressed as means ± SEMs from 3 independent experiments. In all bar diagrams, individual values are shown as small circles. n.s., non-significant; **P ≤ 0.01, *P ≤ 0.05, estimated by two-tailed unpaired Student’s t-test.

### GP63 in *Leishmania* conditioned media targets DICER1 to inhibit maturation of miR-122, a negative regulator of hepcidin

Hepcidin expression in liver has been shown to be negatively regulated by miR-122 (Castoldi et al., 2011). In an unrelated study, it was observed that purified GP63 from *L. donovani* extract cleaves DICER1, an endoribonuclease essential for processing of precursor microRNAs (pre-miRNAs) into its mature and functional form (Ghosh et al., 2013). These prior studies prompted us to check whether treating macrophage cells with Lm-CM could deplete DICER1 levels and, consequently, inhibit the processing of pre-miR-122 in a GP63-dependent manner. As shown in Fig. 6, A and B, we found a significant depletion of DICER1 in macrophage cells treated with Lm-CM or LmCas9/T7-CM, but not in those cells treated with LmGP63^−/-^-CM.

**Figure 6.**
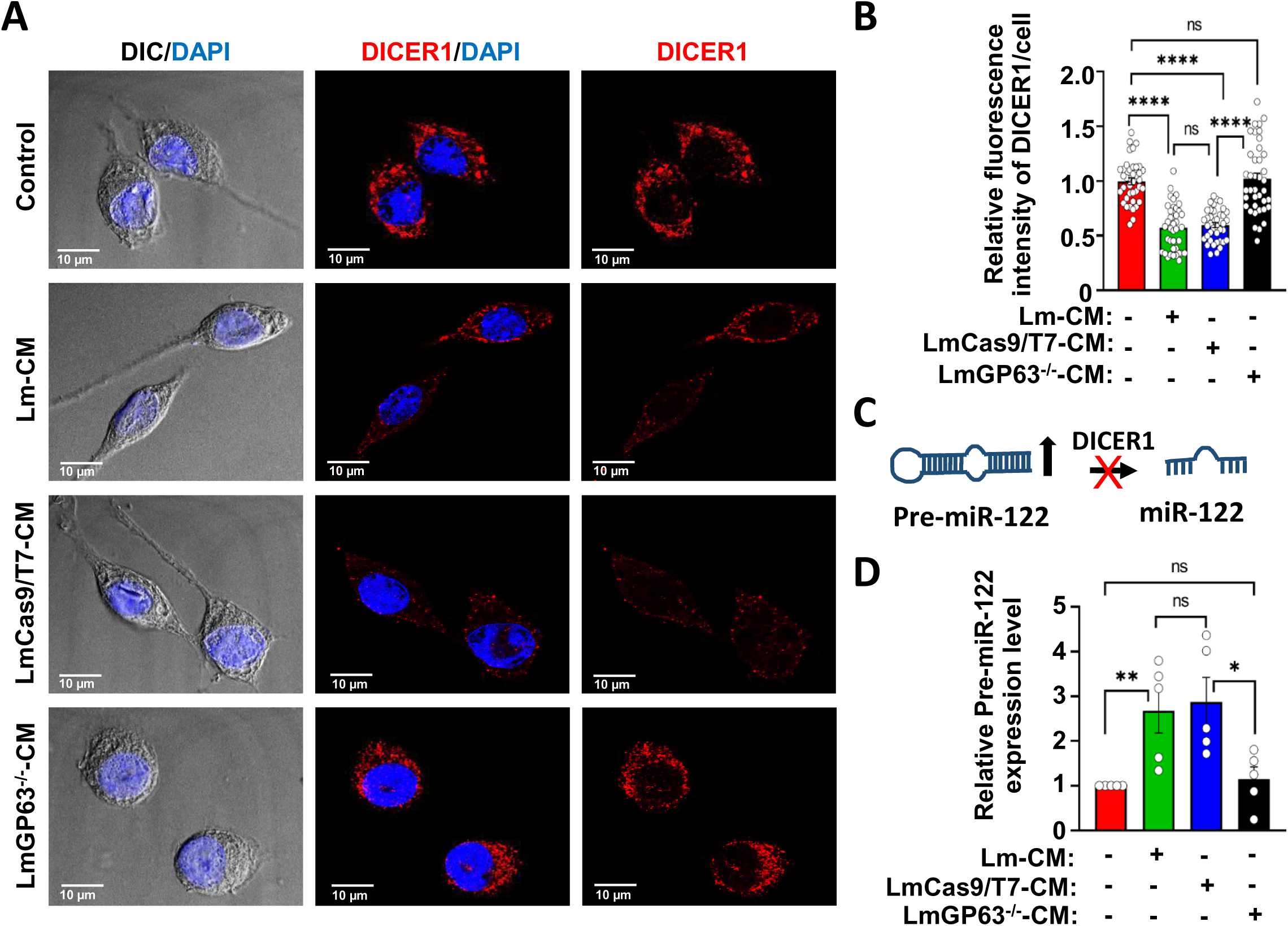
*L. major* GP63 targets DICER1 to inhibit miR-122 maturation in macrophages. **(A)** DICER1 was visualised by immunostaining with anti-DICER1 (red) in J774A.1 macrophages treated for 12 hours with M199 media only (control) or with Lm-CM, LmCas9/T7-CM or LmGP63^−/-^-CM. Nuclei were stained with DAPI (blue) and the DIC/DAPI panel shows the overall cell morphology. Images were acquired with Leica SP8 confocal, 63× objective. **(B)** Quantification of the DICER1 fluorescence intensities in the respective images shown in bar diagram. Values are expressed as means ± SEMs (at least 39 cells from N = 3 independent experiments were analyzed). **(C)** Schematic representation of DICER1-mediated processing of pre-miR-122 to mature miR-122. As shown in the figure, depletion of DICER1 is expected to increase the abundance of pre-miR-122. **(D)** Bar diagram showing qRT-PCR data of relative pre-miR-122 mRNA levels in J774A.1 macrophages treated for 12 hours with M199 media only (control) or with Lm-CM, LmCas9/T7-CM or LmGP63^−/-^-CM. All the measurements were performed using the control cell as reference sample (expression level set to 1.0) and β-actin as an endogenous control gene for normalization. Values are expressed as means ± SEMs from N = 5 independent experiments. In all bar diagrams, individual values are shown as small circles. n.s., non-significant; ****P ≤ 0.0001, **P ≤ 0.01, *P ≤ 0.05 estimated by two-tailed unpaired Student’s t-test.

Since depletion of DICER1 is expected to inhibit the process of microRNA maturation, we checked whether treatment of macrophages with Lm-CM or LmCas9/T7-CM leads to an accumulation of pre-miR-122 due to the processing defect (Fig. 6C). Our qRT-PCR data confirmed that treatment of macrophages with Lm-CM or LmCas9/T7-CM resulted in a significant accumulation of pre-miR-122, indicating a block in miR-122 maturation. However, this impairment was not observed in macrophages treated with LmGP63⁻/⁻-CM, where pre-miR-122 levels remained unchanged (Fig. 6D). Collectively, these results indicate that GP63-mediated depletion of DICER1 is responsible for the inhibition of miR-122 maturation. Given that miR-122 is a negative regulator of hepcidin expression, we propose that GP63 present in Lm-CM induces hepcidin expression in host macrophages by targeting the DICER1/miR-122 axis, thereby facilitating proteasomal degradation of Nramp1.

### GP63 acts through the DICER1/hepcidin axis to lower Nramp1 levels in *Leishmania -* infected BALB/c mice

We next asked whether the mechanism by which GP63 in LM-CM regulates Nramp1 levels in cultured macrophage cells is also operational in an *in vivo* infection model. To investigate this, we infected the footpads of BALB/c mice with wild type *L. major* (LmWT), LmCas9/T7, or LmGP63^−/-^ strains and performed immunofluorescence analysis for Nramp1, DICER1 and hepcidin levels in cryosectioned footpads at 6 and 13 weeks post infection (experiment design illustrated in Fig. 7A). First, we confirmed the presence of GP63 in footpad sections of the mice infected with LmWT or LmCas9/T7 strains and as expected, no GP63 signal was detected in mice infected with the LmGP63^−/-^ strain (Fig. 7B). It may be noted that the mice infected with the LmGP63^−/-^ strain harboured a nearly similar number of parasites in their footpads as those infected with the LmWT or LmCas9/T7 strains at 6 weeks post infection, although their lesion score was slightly lower (Fig. 7, C and D). This data is consistent with a previous report demonstrating that compared to wild type *L. major* infection, GP63-deficient parasites resulted in a mild delay in lesion formation during early infection (∼ 5 weeks), which became more pronounced over time (Joshi et al., 2002). Staining of footpad sections with Nramp1 antibody revealed a significant reduction in Nramp1 levels in mice infected with LmWT or LmCas9/T7 strains at 6 and 13 weeks post infection compared to the uninfected control, whereas the LmGP63^−/-^ strain failed to deplete Nramp1 (Fig. 7, E and F). Co-localization of Nramp1 with CD32 (FcR), a macrophage-specific marker, confirmed that Nramp1 is exclusively expressed in the macrophages of the mouse footpad (Fig. S4). Next, we checked for the DICER1 and hepcidin levels in cryosectioned footpad sections of uninfected BALB/c mice or those infected with LmWT, LmCas9/T7 or LmGP63^−/-^ strains. In agreement with our *in vitro* data, we observed a significant reduction in the DICER1 levels with a concomitant surge of hepcidin levels in the footpad sections of the mice infected with LmWT or LmCas9/T7 strains at 6- and 13-weeks post infection compared to uninfected mice. Footpad sections of the LmGP63^−/-^ mice showed neither a depletion of DICER1 levels nor an increase in hepcidin levels, and these levels were comparable to those in uninfected mice (Fig. 8, A-D). Collectively, these findings provide an unambiguous *in vivo* validation that GP63 indeed acts through the DICER1/hepcidin axis to lower Nramp1 levels in host macrophages.

**Figure 7.**
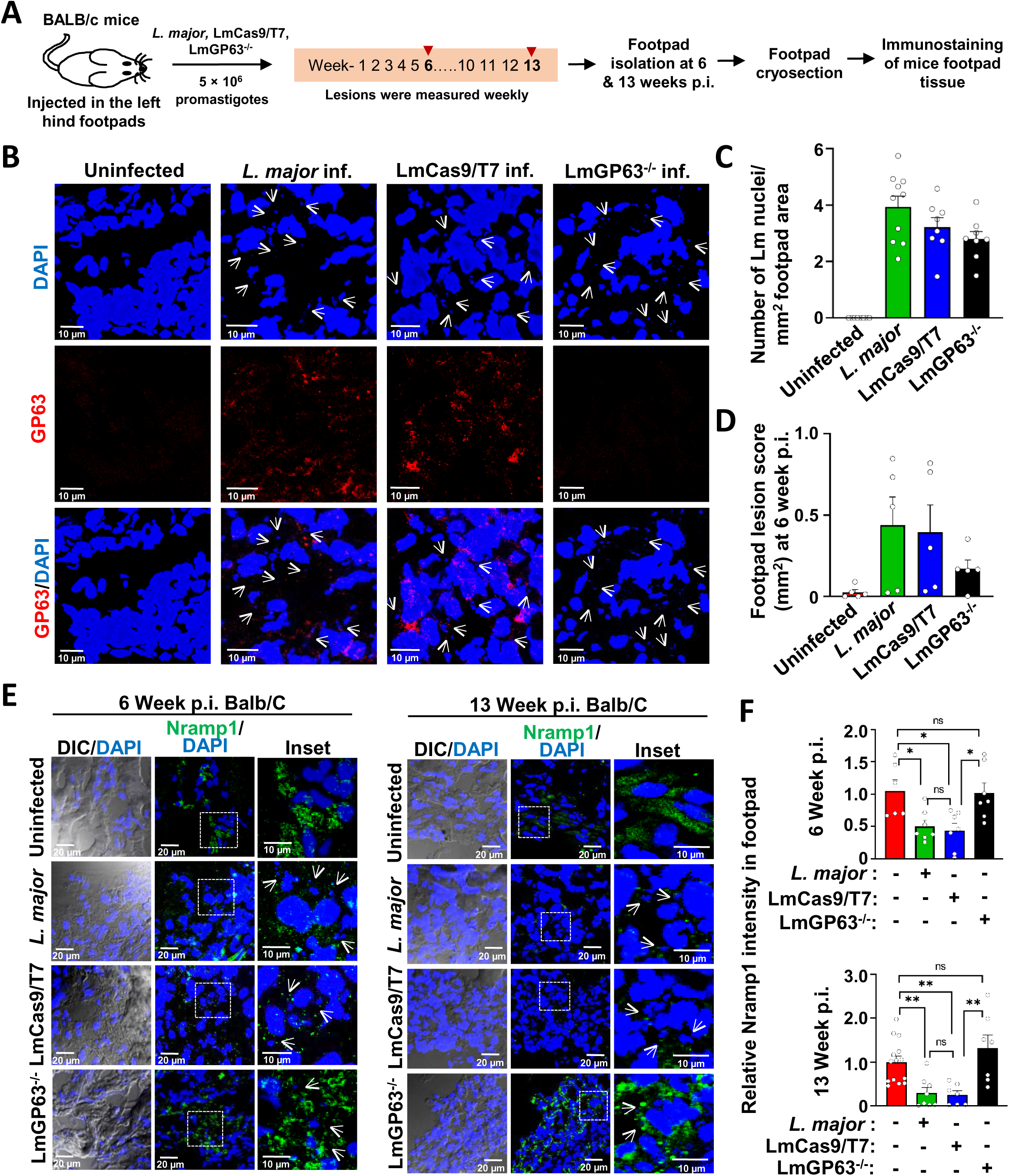
Infection of BALB/c mice with LmGP63^−/-^ parasite failed to deplete Nramp1. **(A)** Schematic illustration of the infection protocol of BALB/c mice with wild type *L. major*, LmCas9/T7, or LmGP63^⁻/⁻^ strains. **(B)** Immunofluorescence staining for GP63 (red) in the footpad cryosections of BALB/c mouse that were either uninfected or infected with wild type *L. major*, LmCas9/T7 or LmGP63^−/-^ strains. Tissues were harvested at 6 weeks post infection (p.i.). Nuclei were stained with DAPI (blue) and small *Leishmania* nuclei are marked with white arrows. **(C)** Bar diagram showing the number of *Leishmania* nuclei per mm² of tissue area quantified by maximum intensity projections (MIP) of the images considering the average nuclear area ∼ 2.5 µm². Individual values are represented as small circles. **(D)** BALB/c mice were infected subcutaneously in the left hind footpad with 5 × 10⁶ stationary-phase promastigotes of wild type *L. major*, LmCas9/T7, or LmGP63⁻/⁻. Footpad swelling was monitored weekly by measuring the width and thickness of the infected foot. Bar diagram showing lesion scores assessed at week 6 p.i. relative to uninfected footpads**. (E)** Immunofluorescence staining for Nramp1 (green) in the footpad cryosections of BALB/c mouse that were either uninfected or infected with wild type *L. major*, LmCas9/T7 or LmGP63^−/-^ strains. Tissues were harvested at 6 (left panel) or 13 (right panel) weeks p.i. Nuclei were stained with DAPI (blue) and small *Leishmania* nuclei are marked with white arrows. **(F)** Quantification of the Nramp1 fluorescence intensities in the respective mice footpad cryosections. Top panel shown the data for 6 weeks p.i. while the 13 weeks p.i. data are in the bottom panel. Values are expressed as means ± SEMs (N = 5 mice). Images were acquired with Leica SP8 confocal, 63× objective. In all bar diagrams, individual values are shown as small circles. n.s., non-significant; **P ≤ 0.01, *P ≤ 0.05, estimated by two-tailed unpaired Student’s t-test.

**Figure 8.**
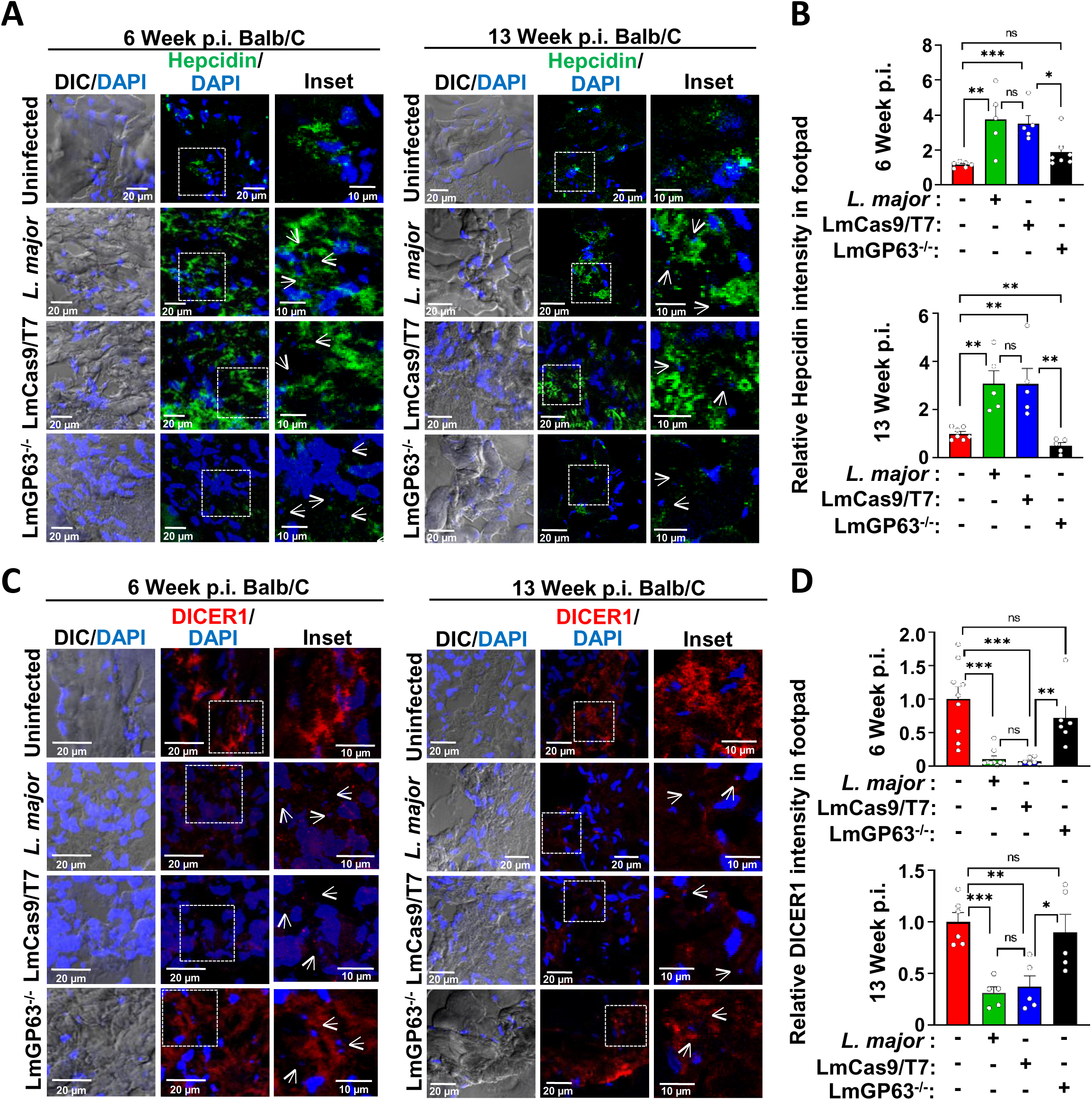
Infection of BALB/c mice with LmGP63^−/-^ parasite failed to upregulate hepcidin expression or deplete DICER1. **(A)** Immunofluorescence staining for hepcidin (green) in the footpad cryosections of BALB/c mouse that were either uninfected or infected with wild type *L. major*, LmCas9/T7 or LmGP63^−/-^ strains. Tissues were harvested at 6 (left panel) or 13 (right panel) weeks p.i. Nuclei were stained with DAPI (blue) and small *Leishmania* nuclei are marked with white arrows. **(B)** Quantification of the hepcidin fluorescence intensities in the respective mice footpad cryosections. Top panel shown the data for 6 weeks p.i. while the 13 weeks p.i. data are in the bottom panel. Values are expressed as means ± SEMs (N = 5 mice). **(C)** Immunofluorescence staining for DICER1 (red) in the footpad cryosections of BALB/c mouse that were either uninfected or infected with wild type *L. major*, LmCas9/T7 or LmGP63^−/-^ strains. Tissues were harvested at 6 (left panel) or 13 (right panel) weeks p.i. Nuclei were stained with DAPI (blue) and small *Leishmania* nuclei are marked with white arrows. **(D)** Quantification of the DICER1 fluorescence intensities in the respective mice footpad cryosections. Top panel shown the data for 6 weeks p.i. while the 13 weeks p.i. data are in the bottom panel. Images were acquired with Leica SP8 confocal, 63× objective. Values are expressed as means ± SEMs (N = 5 mice). In all bar diagrams, individual values are shown as small circles. n.s., non-significant; ***P ≤ 0.001, **P ≤ 0.01, *P ≤ 0.05, estimated by two-tailed unpaired Student’s t-test.

## DISCUSSION

Nutritional immunity is a vital host defense system that deprives pathogens of essential micronutrients, such as iron, to hinder their growth and limit infection (Cassat and Skaar, 2013; Murdoch and Skaar, 2022). A key player in this process is Nramp1, which actively efflux iron from macrophage phagolysosomes, reducing its availability to engulfed pathogens. However, *Leishmania* have evolved a sophisticated strategy to counteract this by causing hepcidin-mediated proteasomal degradation of Nramp1 upon infection. Here, we uncover a crucial role of the parasite-secreted factor GP63 in driving this process. We demonstrate that GP63 targets the DICER1/miR-122 axis in host macrophages to trigger hepcidin expression, ultimately leading to depletion of Nramp1 and enhanced phagolysosomal iron content.

Our quest to identify the parasite-secreted factor responsible for causing Nramp1 depletion was prompted by an interesting observation that Nramp1 levels were diminished not just in *Leishmania* infected cells but also in neighboring uninfected ones. Such unanticipated bystander effect suggested that the parasite’s influence on Nramp1 extended beyond direct infection, possibly through a diffusible factor capable of modulating the host environment. This notion was further fuelled by the fact that a large number of secreted proteins constitute the *Leishmania* exoproteome, and that treatment of macrophages with *L. mexicana*-derived exoproteome has been shown to modulate signalling pathways, including inhibition of nitric oxide production (Hassani et al., 2011). Consistent with our hypothesis, we found that treating macrophages with Lm-CM could cause proteasomal degradation of Nramp1. However, identifying the secretory factor responsible for Nramp1 depletion was a challenge. Multiple studies have shown that the primary mode of protein secretion in *Leishmania* occurs via exosomal vesicles, and only a minor fraction of the secreted proteins contains an N-terminal signal peptide for secretion in free form (Silverman et al., 2010; Pissarra et al., 2022). Interestingly, analysis of the *L. infantum* exoproteome revealed a distinct subset of proteins present in both the exosomal fraction and as free forms, with GP63, a zinc-dependent metalloprotease, being the most abundant among them (Santarém et al., 2013). This prior observation, combined with our data showing that both exosomal and non-exosomal fractions of the Lm-CM retained Nramp1-depleting activity, suggested that GP63 could be the elusive secretory factor driving this effect. We provided definitive evidence for this by exposing macrophages to Lm-CM pre-treated with GP63 inhibitors or to conditioned media from the GP63^−/-^ *L. major*, both of which failed to induce Nramp1 degradation.

Having confirmed that GP63 in Lm-CM is indeed responsible for proteasomal degradation of Nramp1, the next daunting task was to delineate the underlying mechanism. GP63 was originally identified as a major surface glycoprotein of *Leishmania* with Zn-dependent proteolytic activity. It was proposed to play roles in parasite’s attachment to macrophages while also protecting it from phagolysosomal degradation (Russell and Wilhelm, 1986; Chaudhuri et al., 1989). Infection with GP63-deficient parasites resulted in a delayed onset of lesion formation in mice, establishing its role as a virulence factor (Joshi et al., 2002). Subsequent studies confirmed that GP63 is also released into the extracellular space through autoproteolysis at the cell surface as well as via exosomal secretion (McGwire et al., 2002). Interestingly, GP63 was also detected within exosomes released from the *L. mexicana* infected macrophages (Hassani and Olivier, 2013). Exosomal GP63 are rapidly internalized by recipient macrophages via a lipid raft-dependent mechanism and play vital roles in host-pathogen communication (Gomez et al., 2009). Utilizing its proteolytic activity, GP63 is known to alter macrophage signaling and the innate immune response (Isnard et al., 2012). Prior studies have identified protein tyrosine phosphatases (PTPs) as targets of GP63, leading to their activation and subsequent modulation of MAPK and JAK-STAT signaling pathways in macrophages (Gomez et al., 2009). Membrane fusion regulators such as Synaptotagmin XI, VAMP8 and Syntaxin-5 were also reported as host substrates for GP63 (Arango Duque et al., 2014; Matheoud et al., 2013; Guay-Vincent et al., 2022). Purified GP63 from *L. donovani* was shown to cleave the microRNA processing enzyme DICER1, heterologously expressed in HEK293T cells. This DICER1-targeting ability enables *L. donovani* to supress miR-122 in mouse liver, resulting in reduced serum cholesterol levels through downregulation of miR-122-regulated cholesterol biosynthesis genes (Ghosh et al., 2013). A more recent study demonstrated that *L. donovani* GP63 cleaves poly(rC)-binding proteins 1 & 2 (PCB1, and PCB2), which act as chaperones for iron loading into iron storage protein ferritin (Sen et al., 2022). However, GP63 has not yet been shown to directly or indirectly target any iron transport protein of the host cell.

Since Lm-CM-induced depletion of Nramp1 was found to be mediated by proteasomal degradation, the possibility of Nramp1 being a direct target of GP63 was ruled out. This is also supported by our previous finding that *L. major* infection reduces Nramp1 levels through the ubiquitin-proteasome pathway. Earlier, we also demonstrated that *L. major* infection in macrophages led to upregulation of the iron-regulatory peptide hormone hepcidin, which physically interacts with Nramp1. Inhibition of hepcidin transcription prevented Nramp1 degradation, suggesting that Nramp1 depletion in *L. major* infected macrophages is hepcidin-dependent (Banerjee and Datta, 2020). Interestingly, we observed that while treatment of macrophages with Lm-CM significantly enhanced hepcidin expression and Nramp1 ubiquitination, conditioned media from the GP63^−/-^ *L. major* lacked these properties. Together, these findings suggest that GP63 in Lm-CM indirectly promotes Nramp1 depletion by enhancing hepcidin expression.

Hepcidin, primarily produced by the hepatocytes, regulates iron release into the plasma by binding to ferroportin, the iron exporter located on the plasma membrane of target cells, leading to its internalization and degradation (Nemeth et al., 2004). In response to excess iron in circulation, hepatic hepcidin is transcriptionally activated via the BMP6-SMAD pathway. On the hepatocyte surface, Bone Morphogenetic Protein 6 (BMP6) interacts with BMP receptors and the co-receptor hemojuvelin (HJV) to activate SMAD signaling. Phosphorylated SMAD1/5/8 forms a heteromeric complex with SMAD4, which translocates to the nucleus, binds to the *HAMP* (hepcidin) gene promoter, and induces its transcription (Wang et al., 2005; Andriopoulos et al., 2009; Camaschella, 2009). Additionally, hepcidin expression is positively regulated by the hereditary hemochromatosis protein (HFE), which signals through transferrin receptor 2 (TfR2) or BMP type 1 receptor ALK3 (Wu et al., 2014). Interestingly, the microRNA miR-122 has been found to negatively regulate hepcidin production in hepatocytes by inhibiting transcription of *Hjv* and *Hfe*, key activators of hepcidin expression (Castoldi et al., 2011). These previous reports on hepcidin regulation in hepatic cells prompted us to ask whether similar regulatory circuit exists in macrophages and if *L. major* GP63 can tweak it to stimulate hepcidin expression. Since, microRNA-processing enzyme DICER1 has been identified as a substrate for *L. donovani* GP63, we were curious to examine the status of DICER1 and miR-122 in macrophage cells treated with Lm-CM or conditioned media from the GP63^−/-^ *L. major* (Ghosh et al., 2013). Strikingly, treatment with Lm-CM, but not with conditioned media from the GP63^−/-^ parasite, caused significant depletion of DICER1 and impaired processing of pre-miR-122 to mature miR-122 in macrophages. Given that miR-122 is a known repressor of hepcidin, the GP63-mediated processing defect of miR-122 provides a mechanistic explanation for the upregulation of hepcidin in Lm-CM-treated macrophages and the consequent hepcidin-mediated proteasomal degradation of Nramp1 (Castoldi et al., 2011). Does the mechanism by which GP63 in LM-CM targets DICER1/miR-122/hepcidin axis to deplete Nramp1 levels in cultured macrophage cells also operates under actual infection conditions? This was tested in an animal model of *L. major* infection in BALB/c mice. Compared to wild type *L. major*, the GP63 deficient parasite (LmGP63^−/-^) clearly lacked the ability to deplete DICER1, induce hepcidin expression or reduce Nramp1 levels in the infected mouse footpad, providing *in vivo* validation for our *in vitro* observations.

To summarize, we uncover a previously unrecognized regulatory axis controlling Nramp1 levels in macrophages, which is hijacked by *Leishmania* GP63 to sequester iron within phagolysosomal compartment, thereby supporting parasite survival (Fig. 9). It remains to be seen if this miR-122/hepcidin-mediated post-translational regulation of Nramp1 have other physiological implications beyond influencing the infection outcome. The role of Nramp1 in facilitating efficient iron recycling during erythrophagocytosis is well-established (Soe-Lin et al., 2008, 2009). Notably, overexpression of Nramp1 has been shown to elevate redox-active labile iron pool in the cytosol, likely due to enhanced iron export from the phagolysosomal compartment (Soe-Lin et al., 2008). The labile iron pool being a potent inducer of oxidative stress and cell death (Dixon and Stockwell, 2014), it would be intriguing to explore whether miR-122/hepcidin-mediated depletion of Nramp1 serves as a protective mechanism to mitigate iron-induced cytotoxicity, particularly during episodes of heightened erythrophagocytosis.

**Figure 9.**
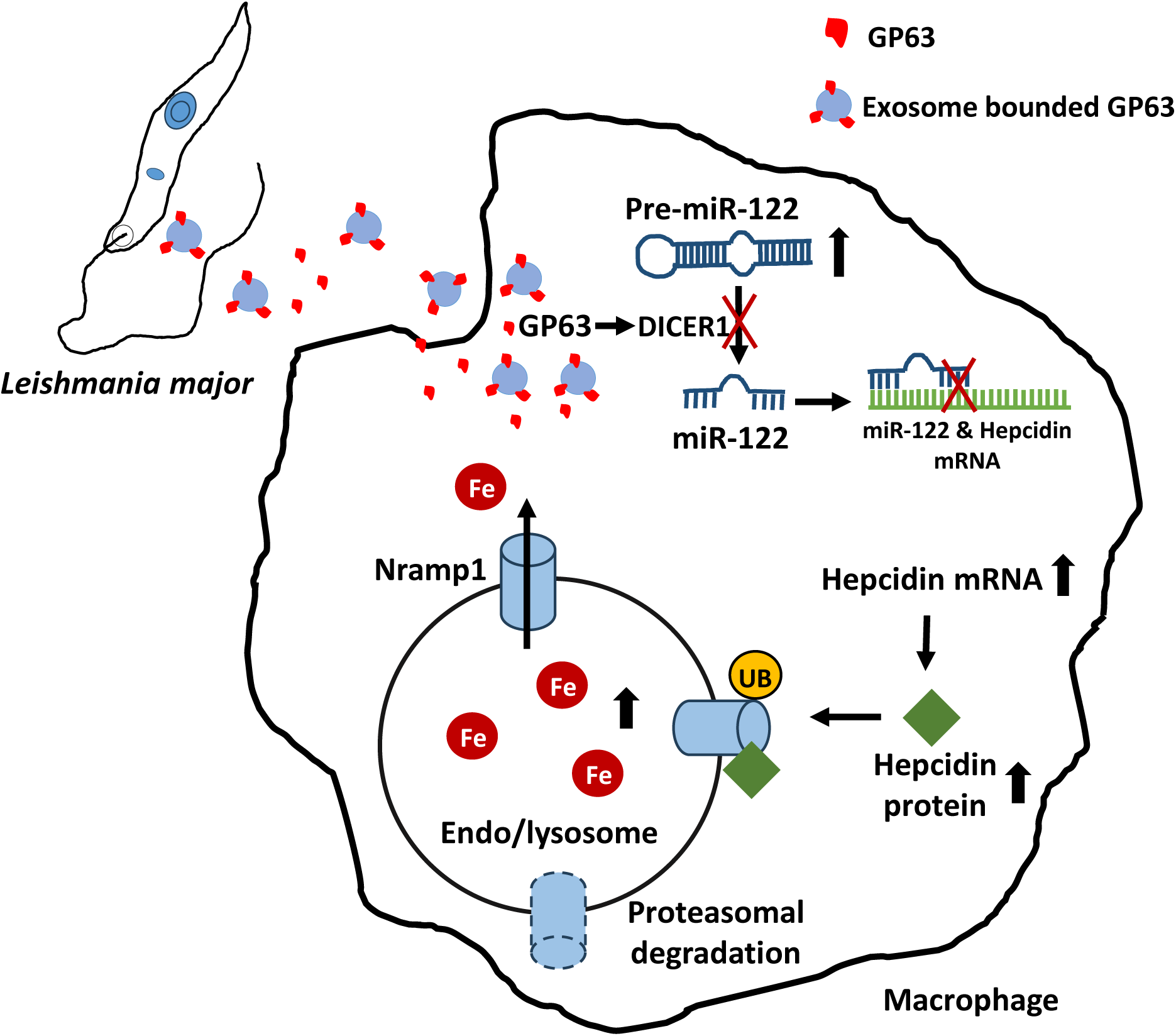
Schematic representation of GP63-mediated Nramp1 degradation mechanism. *Leishmania*-secreted GP63 targets DICER1 in host macrophages, blocking maturation of miRNA-122, a known negative regulator of hepcidin. This triggers hepcidin expression, ultimately leading to hepcidin-mediated proteasomal degradation of Nramp1 and enhanced endo/lysosomal iron content.

## MATERIALS AND METHODS

Reagents, antibodies and primers/plasmids used in this study are listed in Tables S1, S2 and S3, respectively. Unless otherwise specified, all other reagents were obtained from Sigma-Aldrich.

### *L. major* culture and preparation of conditioned medium

*L. major* promastigotes (strain 5ASKH, kindly gifted by Dr. Subrata Adak, CSIR-IICB, Kolkata) were grown at 26^0^C in M199 medium supplemented with 23.5 mM HEPES, 10 µg/ml hemin, 150 µg/ml folic acid, 0.2 mM adenine, 120 U/ml penicillin, 60 µg/ml gentamycin, 120 µg/ml streptomycin and 15% heat-inactivated FBS (Pal et al., 2017). Cas9 and T7 RNA polymerase expressing *L. major* cells were grown similarly with additional supplementation of 50 μg/ml hygromycin. The GP63^−/-^ *L. major* strain was maintained with additional supplementation of 20 µg/ml puromycin and 5 µg/ml blasticidin.

The conditioned media from *L. major* promastigotes (wild type or the engineered strains) were prepared from late log phase culture (3.6 × 10^7^ cells/mL) of parasites. The culture was centrifuged at 1200 x g for 10 min at 26^0^C to obtain cell free supernatant, which was further centrifuged at 10,000 x g for 30 minutes to remove any debris. The conditioned media, containing secretory factors, was sterilized by passing through 0.22 μm filter units. The absence of intact parasites or their cytosolic content in the collected conditioned media was confirmed by microscopic observation and western blot using antibodies against known cytosolic proteins of *Leishmania*. Wherever mentioned, the conditioned media was heated at 95°C for 5 minutes and then cooled to room temperature before being used to treat macrophages. For protease digestion, trypsin was added to the conditioned media at a final concentration of 0.25%, mixed thoroughly, and incubated at 37°C for 10 minutes. Thereafter, a protease inhibitor cocktail was added to inactivate the trypsin prior to using it for treatment of macrophages. For pharmacological inhibition of GP63, the conditioned media was incubated with divalent metal chelators EDTA (1 mM) or 1, 10-Phenanthroline (1 mM) at 37°C for 2 hours.

### Infection of macrophages with *L. major* and treatment with conditioned medium

J774A.1 murine macrophage cell line (ATCC #TIB-67) were cultured in Dulbecco’s Modified Eagle Medium (DMEM) at 37°C in a humidified CO_2_ (5%) incubator. The medium was supplemented with 2 mM L-glutamine, 100 U/ml penicillin, 100 μg/ml streptomycin, and 10% heat-inactivated FBS, and adjusted to pH 7.4. Following our previously published protocol, the LPS-activated macrophages were infected with *L. major* at a ratio of 1:30 (Pal et al., 2017). After 12 hours of infection, cells were washed with PBS and processed for immunofluorescence.

Uninfected macrophages (∼6 hours post seeding) was treated for 12 hours with *L. major* conditioned media [100ul or 1ml conditioned media for 1.2 × 10^5^ (for immunofluorescence studies) or 1.2 × 10^6^ (for western blot experiments) macrophage cells, respectively].

### Isolation of peritoneal macrophage from BALB/c mice

Thioglycolate-elicited peritoneal macrophages were isolated from 6 to 8-week-old BALB/c mice, as described earlier (Zhang et al., 2008). The mice were maintained at the IISER Kolkata animal facility following CCSEA guidelines and the experiment was conducted as per Institutional Animal Ethics Committee (IAEC) approved protocol After four days of intraperitoneal injection with 3% Brewer’s thioglycolate medium, mice were euthanized, and peritoneal macrophages were extracted using a 20G needle. The collected macrophages were then cultured and treated with *L. major* conditioned media as described above.

### Isolation of exosomal vesicles from *L. major* conditioned medium

The exosomal vesicles (Lm-EVs) were isolated from the *L. major* conditioned media using an exosome isolation kit following the protocol suggested by the manufacturer. Briefly, 0.5 ml of the total exosome isolation reagent was added to 1ml of the conditioned media and mixed well by pipetting. Mixture was then incubated overnight at 4^0^C and centrifuged at 10,000 x g for 1 hour at 4^0^C to pellet the Lm-EVs. The Lm-EVs were resuspended in appropriate volume of M199 media or PBS. Alongside, the post-EV supernatant was also collected. Both the fractions (Lm-EVs and post-EV supernatant) were used for the treating the macrophages or SEM.

### Scanning electron microscopy of purified extracellular vesicles

For scanning electron microscopy (SEM), purified extracellular vesicles (EVs) were isolated from *L. major* CM. The EVs were diluted with filtered PBS, and 10 μL of the sample was deposited on a mica sheet, and then incubated at room temperature until dry. The sample was washed with PBS and fixed with 2.5% glutaraldehyde for 40 minutes at room temperature. Following fixation, the EVs were washed with PBS and further fixed with OsO₄ for 5 minutes at room temperature. The EVs were washed twice with PBS and then with Milli-Q water to remove any residual salts. Finally, the EVs were visualized using a Zeiss Supra 55VP scanning electron microscope (SEM).

### Generation of GP63 knockout *L. major* strain (LmGP63^−/-^) by CRISPR/Cas9

LmGP63^−/-^ strain was generated on the background of an *L. major* strain stably expressing Cas9 and T7 RNA polymerase (LmCas9/T7) that was recently developed by us (Seth et al., 2025). At first, the LmCas9/T7 strain was verified by genomic DNA PCR with Cas9 FP/Cas9 RP and T7 pol FP/T7 pol RP primer sets for amplification of Cas9, T7 RNA polymerase, respectively. The rRNA45 gene, amplified with rRNA45 FP/ rRNA45 RP primer set, was used as control (Fig. S3A). We planned to knock out both the alleles of all four copies of GP63 following the method described by Beneke *et al*. and as illustrated in Fig. 4 A and B (Beneke et al., 2017). Briefly, 5’ GP63sgRNA and 3’ GP63sgRNA oligonucleotides were used to generate sgRNA templates. These oligonucleotides contained the T7 promoter, sgRNA sequence for specifically targeting 20-nucleotide long sequences at 354 bp upstream of 5’ UTR of GP63-1 ORF or 512 bp downstream of 3’ UTR of GP63-4 ORF and a complementary sequence for the sgRNA scaffold. To amplify the sgRNA templates, PCR reactions were set up using G00 primer (which provides the sgRNA scaffold) and 5’ GP63sgRNA or 3’ GP63sgRNA oligonucleotides. Simultaneously, the puromycin and blasticidin resistance cassettes containing 30-nt overhangs of 5’ UTR of GP63-1 ORF and 3’ UTR of GP63-4 ORF were PCR-amplified using pTBLAST or pTPURO plasmids as templates and GP63-1 5’UTR and GP63-1 3’UTR primer sets. All four PCR products (5’ GP63sgRNA template, 3’ GP63sgRNA template, puromycin and blasticidin resistance cassettes) were co-transfected into mid-log phase LmCas9/T7 promastigotes by electroporation. Prior to electroporation, the PCR products were heat-sterilized at 94_°_C for 5 minutes. The LmGP63^−/-^ strain, generated upon integration of the puromycin and blasticidin resistance cassettes into the genome at the sgRNA target sites replacing the GP63 ORFs, was selected in presence of 20 μg/ml puromycin and 5 μg/ml blasticidin. The authenticity of the strain was confirmed by various methods as described in the results section.

### Cell lysis and western blot

J774A.1 macrophages were scrapped from tissue culture plates and collected in microfuge tubes whereas suspension culture of *L. major* was taken directly in the microfuge tubes. The cells were then pelleted at 1200g for 5 minutes, washed twice with ice-cold PBS and then dissolved in lysis buffer (PBS, 1x EDTA-free protease inhibitor cocktail and 2 mM PMSF). The cells were lysed by sonication to prepare the whole cell lysates (WCL). Soluble proteins from the *L. major* conditioned medium were precipitated using chilled acetone, resuspended in PBS, and denatured in Laemmli buffer. The WCL (macrophage or *L. major*) or suspension of conditioned medium were resolved by SDS gel electrophoresis and transferred to activated PVDF membrane. The membrane was blocked in 5% skim milk in 0.05% TBST and probed primary antibodies (rabbit anti-Nramp1, 1:1000; rabbit anti-γ-actin, 1:4000; rabbit anti-hepcidin, 1: 500; mouse anti-Dicer1, 1:1000; rabbit anti-Ldactin, 1:5000; rabbit anti-tubulin antibody, 1:4000 and rabbit anti-LmPGAM, 1:2000. Following washing, blots were then probed with appropriate HRP-conjugated secondary antibody (goat anti-rabbit, 1:4000 or rabbit anti-mouse, 1:4000). The blots were developed using Super Signal West Pico Chemiluminescent Substrate and bands intensities were quantified using ImageJ software, with γ-actin as the loading control.

### Immunofluorescence microscopy

For immunofluorescence studies, macrophages were seeded in 6-well tissue culture plates containing 22 mm coverslips at a density of 1.2 × 10^5^ for 6 hours. For, *L. major* promastigotes, log phase cells were directly mounted on a poly-L-lysin coated glass coverslip. The cells were fixed with acetone: methanol (1:1) for 10 mins at room temperature and permeabilized with 0.1% tritonX -100. After two PBS washes, cells were blocked with 0.2% gelatin for 2 hours at room temperature. Cells were washed with PBS and incubated with primary antibodies (rabbit anti-Nramp1, 1:100; mouse anti-GP63, 1:600; rabbit anti-Hepcidin, 1: 100; mouse anti-Dicer1 1:100) for 1 hour at room temperature. Following another PBS wash, cells were incubated with secondary antibodies (goat anti-rabbit Alexa Fluor 488, 1:800; goat anti-mouse Alexa Fluor 568, 1:800) for 2 hours at room temperature. Cells were again washed with PBS and mounted with an antifade mounting media containing DAPI. Images were acquired with Leica SP8 confocal, Carl Zeiss Apotome.2 or Olympus IX-81 epifluorescence microscope using oil immersion 63× or 40x objectives. Relative fluorescence intensities were measured from Z planes that were merged as a maximum intensity projection (MIP) where ROI was selected for individual cells using the microscope’s software ZEN Blue.

### Flow cytometry

GP63 surface expression in *L. major* promastigotes was analyzed by flow cytometry as previously described (Joshi et al., 2002). Briefly, *L. major* promastigotes (5 × 10⁶ cells) were harvested and washed twice with ice-cold PBS containing 0.5% BSA. The cells were labelled with mouse anti-GP63 primary antibody (1:600) and incubated at 4°C for 1 hour. After primary antibody incubation, the cells were washed twice with containing 0.5% BSA and incubated with goat anti-mouse Alexa Fluor 568 secondary antibody (1:800) at 4°C for 1 hour. The cells were washed again with 1x PBS containing 0.5% BSA and resuspended in 1ml 1x PBS. GP63 expression was analyzed using a BD FACSVerse flow cytometer.

### Gelatin zymography assay

The activity of GP63 in *L. major* whole cell lysate (WCL) and conditioned medium was measured by gelatin zymography assay (Hassani et al., 2014). *L. major* promastigotes (1×10^7^) were washed twice with ice-cold 1x PBS and resuspended in lysis buffer containing 1x PBS, EDTA-free protease inhibitor cocktail, 2mM PMSF. The cells were sonicated to prepare the WCL. *L. major* WCL or conditioned medium were mixed with loading dye containing glycerol and bromophenol blue and both samples were resolved on 10% SDA-PAGE gel copolymerized with 0.12% fish gelatin. Following electrophoresis, the resolved gel was incubated in a buffer (pH 7.4) containing 50 mM Tris, 5 mM CaCl_2_, 2.5% Triton X and 1 μM ZnCl_2_ for 1 hours at room temperature. After incubation, the gels were rinsed in distilled H_2_O, and GP63 activity was detected by incubating overnight in reaction buffer (pH 7.4) containing 50 mM Tris, 5 mM CaCl_2,_ and 1 μM ZnCl_2_ at 37^0^C. The gels were visualized by staining with Coomassie blue and subsequently destained to reveal the zones of proteolytic activity.

### Iron quantification in the endo/lysosomal compartments

Endo/lysosomal vesicles were isolated from the macrophage cells using sucrose density gradient centrifugation as described by us previously (Banerjee and Datta, 2020). Briefly, macrophages were washed with ice-cold 1X PBS and resuspended in lysis buffer containing 1X PBS, EDTA-free protease inhibitor cocktail, and 2mM PMSF. The cell lysate was prepared by passing the suspension through a 26G needle several times, followed by centrifugation at 3,000 rpm for 5 minutes to remove nuclei and unbroken cells. The resulting supernatant was then centrifuged at 12,000 rpm for 6 minutes. After separating the supernatant in the sucrose gradient, the phagosome-enriched 4% sucrose fraction was collected. The iron content (Fe²⁺) of the phagosomal fraction was quantified using a ferrozine assay (Banerjee and Datta, 2020). The Endo/lysosomal fraction was incubated with iron-releasing reagent (10mM HCL and 4.5% KMnO4) for 2 hours at 60^0^ C. After cooling the samples to room temperature, 30μl iron detection reagent (6.5 mM neocuproine, 6.5 mM ferrozine, 2.5 M ammonium acetate, and 1 M ascorbic acid) was added. The mixture was incubated for 30 minutes, and the absorbance was measured at 550 nm using a microplate reader. A standard curve was prepared using a different standard concentration of FeCl_3_ (0-300 μM), and sample iron concentration of samples was calculated from the FeCl_3_ standard curve.

### RNA isolation and quantitative RT-PCR (qRT-PCR)

Total RNA was isolated from J774A.1 macrophages using TRIzol reagent. The isolated RNA was treated with DNase1 to remove any DNA contamination. Using Verso cDNA synthesis kit to cDNA were synthesized from 1μg of the total RNA. Real-time quantitative PCR was performed on 7,500 Real-time PCR systems (Applied Biosystems) using SYBR green fluorophore. Transcript levels of Nramp1, Pre-miR-122, and hepcidin were quantified using respective primers. The transcript levels were normalized with respect to β-actin gene expression. The relative expression levels were calculated using the comparative Ct method (Livak and Schmittgen, 2001).

### Infection of BALB/c mice with *L. major*

As described by us previously, infection experiments in BALB/c mice were performed at IISER Kolkata animal facility following IAEC-approved protocol (Banerjee and Datta, 2023). Briefly, 6 to 8 weeks old female BALB/c mice were subcutaneously injected in the left hind footpad with 5×10^6^ late stationary-phase *L. major* promastigotes (five mice/group in each of the experimental set). As uninfected control, the mice were injected with PBS. Lesion development was assessed weekly by measuring the width and thickness of the infected footpad using a digital calliper. As an internal control, the same measurements were taken from the uninfected footpad of each mouse. The lesion score was determined using the following formula: (width of infected footpad - width of noninfected footpad) × (thickness of infected footpad - thickness of noninfected footpad).

### Cryosectioning and immunostaining of mice footpads tissue

At 6-week and 13-week post infected mice were euthanized and infected mice footpads were harvested and immediately to embed in optimal cutting temperature (OCT) compound. The embedded tissues were then kept at -20°C overnight to ensure complete freezing. 5 µm thick cryosections were cut using Leica CM1950 cryostat and sections were taken on the poly-L-lysine coated glass slides. Sections were air-dried and fixed immediately with 4% PFA for 10 minutes, followed by washing with PBS. Tissue sections were then permeabilized with 0.2% triton X-100 in PBS for 15 minutes. After washing with PBS (2 times) sections were blocked with 0.2% gelatin in PBS for 30 minutes and incubated with primary antibodies (rabbit anti-Nramp1, 1:100; mouse anti-GP63, 1:500; rabbit anti-Hepcidin, 1: 100; mouse anti-Dicer1,1:100; rat anti-FcƔ, 1:100) for 1 hour. Sections were then washed with PBS (3 times) and incubated with secondary antibodies for 1.5 hours at room temperature. Tissues were finally mounted using Vectashield mounting media with DAPI and imaged in Leica SP8 confocal microscope.

### Statistical analysis

All statistical analyses were performed using the Student’s t-test, and the results were expressed as the mean ± SEM from at least three independent experiments. Levels of statistical significance were indicated as follows: a p-value of less than 0.05 was considered statistically significant, with significance levels denoted as *p ≤ .05, ** p < .01, *** p < .001, ****p < .0001.

## ACKNOWLEDGEMENTS

This work was supported by ICMR research grant No: 6/9-7(318)/2023-ECD-II and West Bengal DSTBT grant No: 398(Sanc.)/STBT-13015/16/2024-ST SEC awarded to RD. The authors thank IISER Kolkata central facility staff members: Mr. Anu Das for assistance with animal experiments, Mr. Tamal Ghosh for FACS analysis and Mr. Kashinath Sahu for SEM imaging. We are grateful to Drs. Subrata Adak, Amogh Sahasrabuddhe, Frédéric Bringaud and William S. Sly for providing plasmids for CRISPR-Cas9, LdActin antibody, LmPGAM antibody and hepcidin antibody, respectively. We also thank Drs. Sankar Maiti, and Jayasri DasSarma for sharing critical reagents and for their valuable suggestions. Special thanks to Dr. Budhaditya Mukherjee for providing access to the cryosectioning facility, and to Dr. Subhankar Dolai for his critical comments on the manuscript. SS was supported by a UGC-NET fellowship, Government of India.

## AUTHOR CONTRIBUTION

S.S., and R.D. conceptualized and designed the study. S.S. performed the majority of the experiments, while S.B. contributed to some initial experiments. S.S. analyzed the data, prepared the figures, and wrote the initial draft of the manuscript. S.B. also contributed to manuscript writing. R.D. supervised the work, acquired funding, and wrote the final version of the manuscript.

## Supplementary information

**Figure S1.**
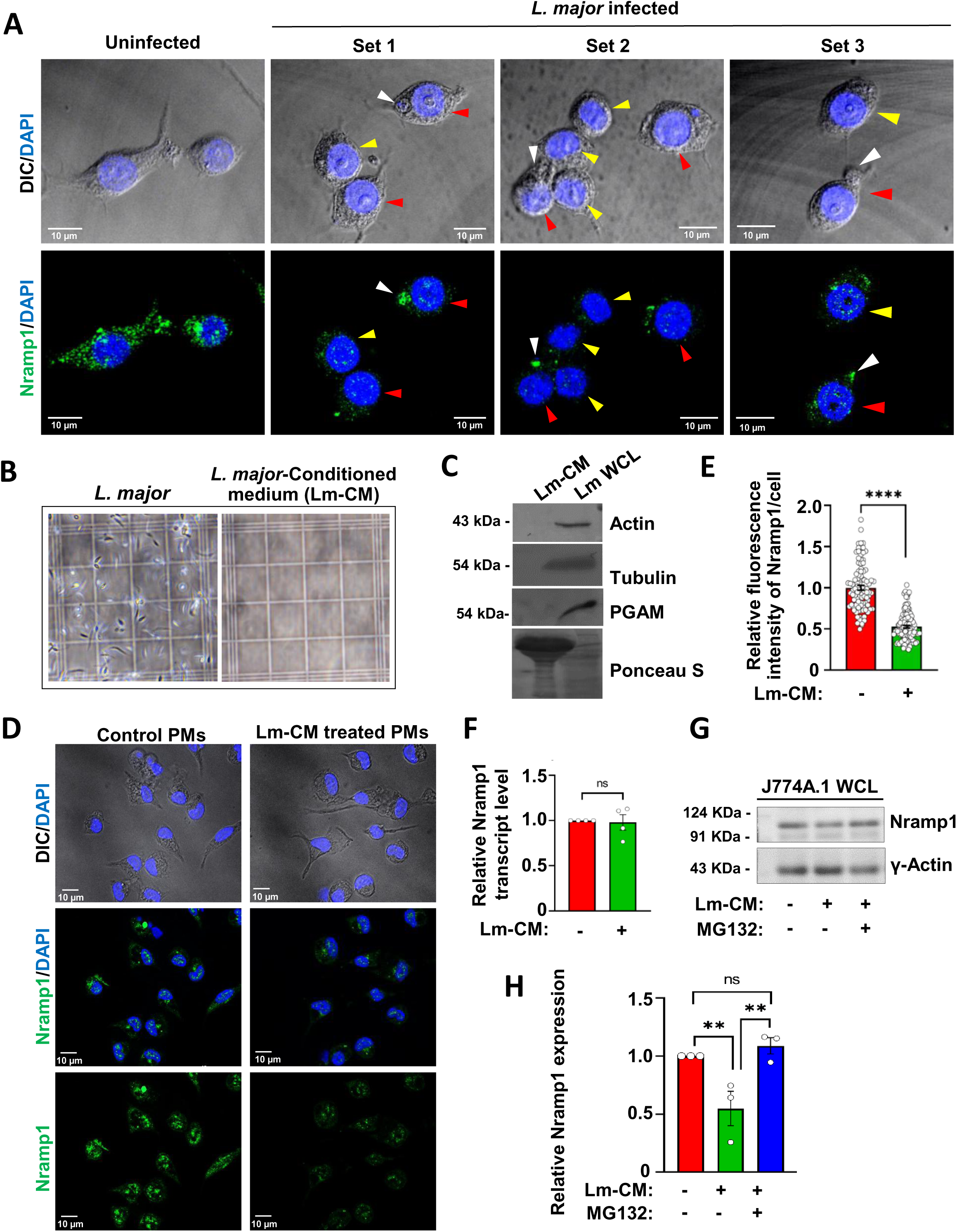
Infection with *L. major* or treatment with *L. major* conditioned medium induces Nramp1 degradation in peritoneal macrophages. **(A)** Nramp1 was visualised by immunostaining with anti-Nramp1 (green) in uninfected or *L. major* (Lm)-infected J774A.1 macrophages (3 fields from different experimental sets). Nuclei were stained with DAPI (blue). DIC/DAPI panel shows the presence of intracellular parasites (smaller nuclei, indicated by white arrows) in infected cells In all the three sets of Lm-infected macrophages, decrease in Nramp1 fluorescence intensities were seen in the the infected cells (marked by red arrows) as well as in bystander uninfected cells marked by yellow arrows). Images were acquired with Leica SP8 confocal, 63× objective **(B)** Light microscope images of *L. major* promastigotes and *Leishmania* condition medium (Lm-CM) confirming the absence of any parasite in it. **(C)** Western blot with antibodies against cytosolic proteins of *Leishmania* (actin, tubulin, and PGAM ) on Lm-CM or *L. major* whole cell lysates (Lm WCL). Ponceau S stained membrane shows total proteins in Lm-CM and Lm WCL. For Lm WCL, 15mg protein was loaded whereas for Lm-CM ∼ 50mg protein was loaded to confirm the absence of any cytosolic proteins of the parasite in Lm-CM. **(D)** Nramp1 was visualised by immunostaining with anti-Nramp1 (green) in BALB/c mice derived peritoneal macrophages treated for 12 hours with *L. major* conditioned medium (Lm-CM) or with M199 medium only (control). Nuclei were stained with DAPI (blue) and the DIC/DAPI panel shows the overall cell morphology. Images were acquired with Carl Zeiss Apotome.2 microscope, 63× objective. **(E)** Quantification of the Nramp1 fluorescence intensities in the respective images shown in bar diagram. The data are expressed as means ± SEM (at least 99 cells from N = 3 independent experiments were analyzed). **(F)** Bar diagram showing qRT-PCR data of relative Nramp1 expression in J774A.1 macrophages treated for 12 hours with M199 media only (-) or with Lm-CM (+). The measurements were performed using the untreated cell as reference sample (expression level set to 1.0) and β-actin as an endogenous control gene for normalization. Values are expressed as means ± SEMs from N = 4 independent experiments. **(G)** Representative western blots of Nramp1 and Ɣ-actin (loading control) on J774A.1 macrophage whole cell lysates (WCL) prepared from either cells treated with just with M199 media (-) or with Lm-CM or Lm-CM + 1µm MG132 (macrophages were pre-treated with MG132 prior to Lm-CM treatment). **(H)** Bar diagram showing the quantification of Nramp1 band densities in the respective samples normalized to Ɣ-actin. Values expressed as means ± SEMs from at least three independent experiments. In all bar diagrams, individual values are shown as small circles. n.s., non-significant; ****P ≤ 0.0001,**P ≤ 0.01estimated by two-tailed unpaired Student’s t-test.

**Figure S2.**
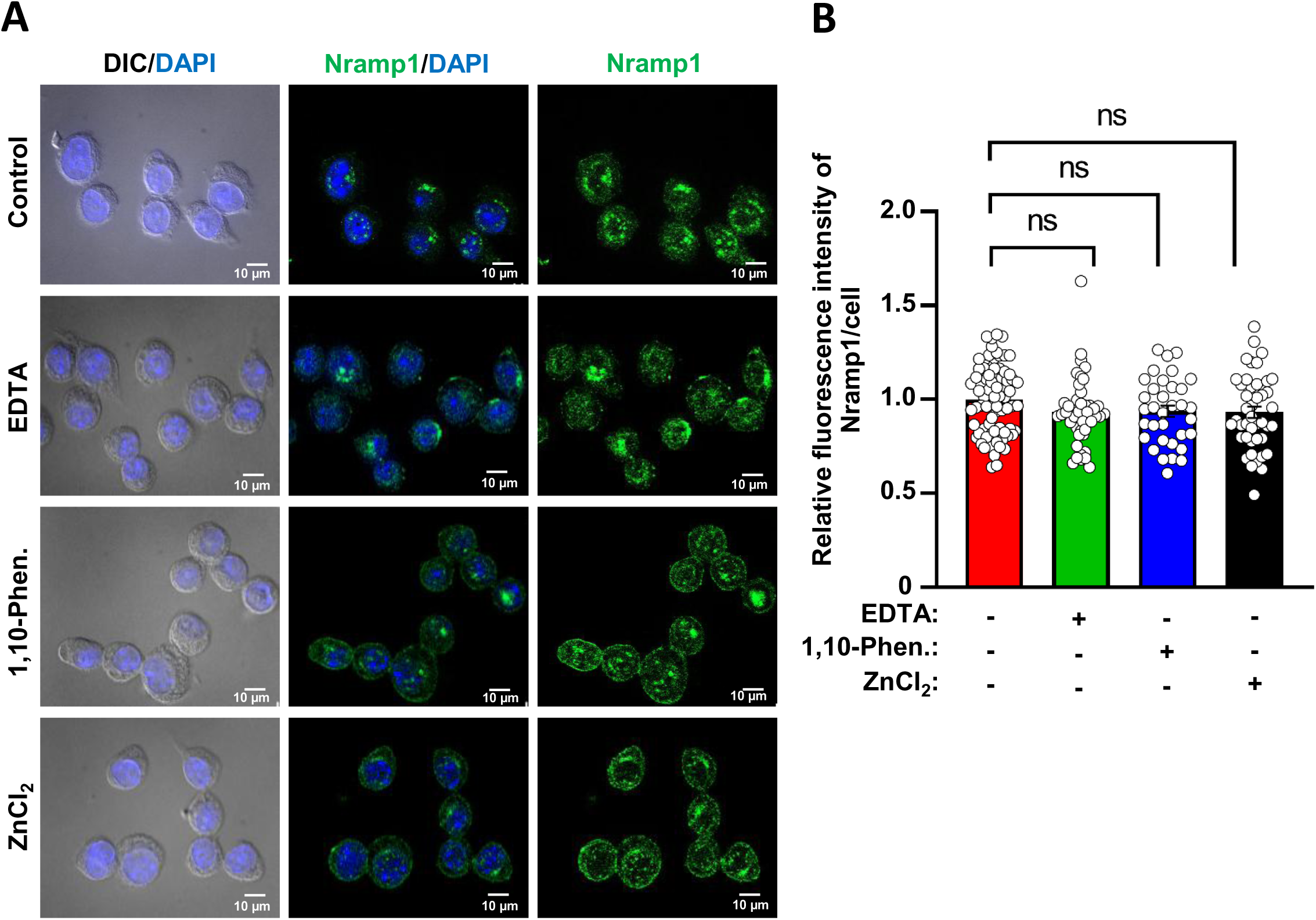
Effect of EDTA, 1,10-Phenanthroline and ZnCl_2_ on Nramp1 expression. **(A)** Nramp1 immunostaining (green) in J774A.1 macrophages treated for 12 hours with just M199 media (control) or M199 + 1mM EDTA, M199 + 1mM 1,10-Phenanthroline or M199 + 1mM ZnCl_2_. Nuclei were stained with DAPI (blue) and the DIC/DAPI panel shows the overall cell morphology. Images were acquired with Carl Zeiss Apotome.2 microscope, 63× objective. **(D)** Quantification of the Nramp1 fluorescence intensities in the respective images shown in bar diagram. Values are expressed as means ± SEMs (at least 35 cells from N = 3 independent experiments were analyzed). In the bar diagram, individual values are shown as small circles. n.s., non-significant, estimated by two-tailed unpaired Student’s t-test.

**Figure S3.**
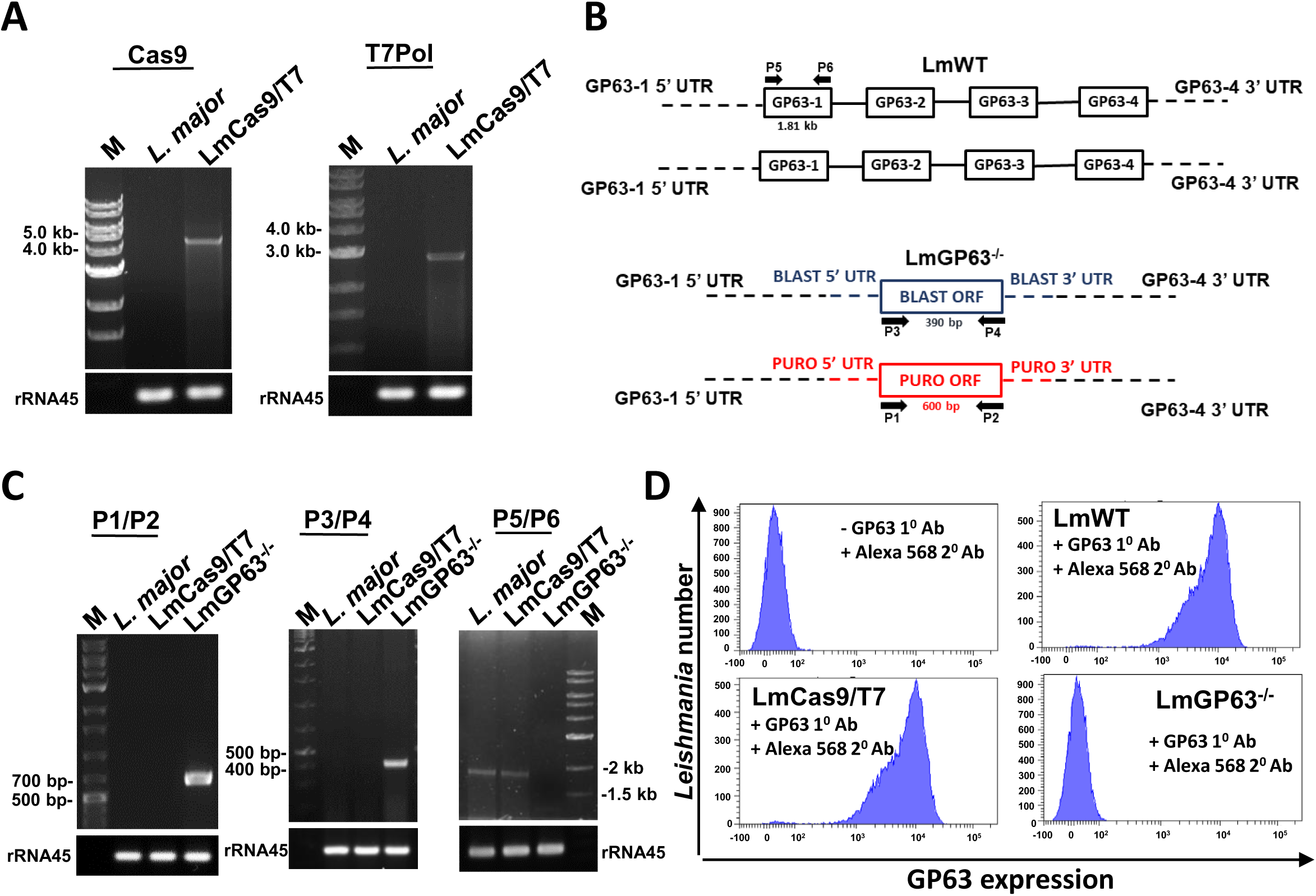
Verification of the LmCas9/T7 and LmGP63^−/-^ strains. **(A)** Agarose gel images showing PCR amplification of the Cas9 and T7 polymerase genes (using gene-specific primers) using LmCas9/T7 genomic DNA as template but not the wild type *L. major* genomic DNA template. **(B)** Schematic representation of the organization of GP63 genes in *L. major* genome. Primers used for the verification of the LmGP63^−/-^ strain are shown. **(C)** Agarose gel images showing genomic DNA PCR results confirming the presence of puromycin (600 bp product with primers P1/P2) and blasticidin (390 bp product with primers P3/P4) cassettes and absence of the GP63 gene (checked with primers P5/P6) in the LmGP63^−/-^ strain. **(D)** FACS analysis of with only Alexa-568-conjugated anti-mouse antibody or mouse anti-GP63 + Alexa-568-conjugated anti-mouse antibody confirming the expression of GP63 in wild type L. major and the LmCas9/T7 strain but not in the LmGP63^−/-^ strain.

**Figure S4.**
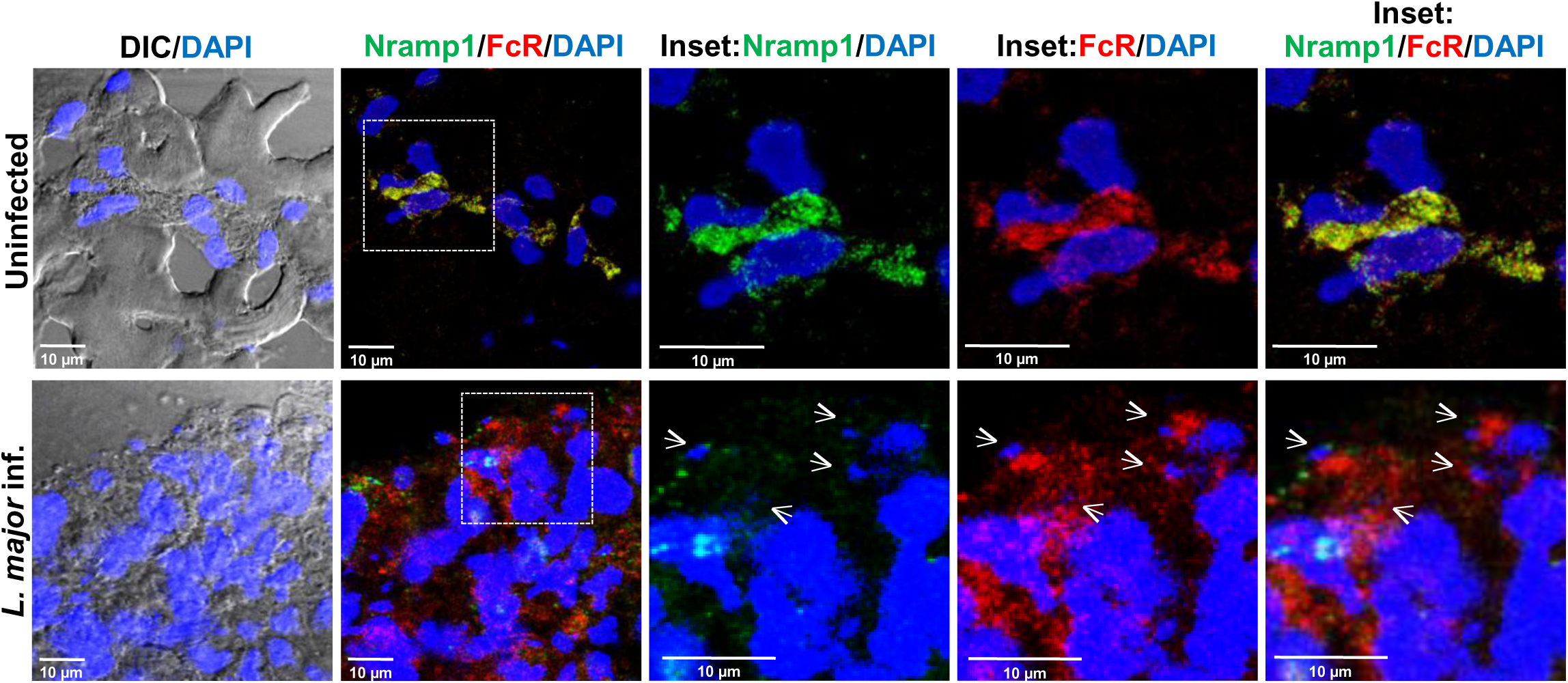
Co-localization of Nramp1 and CD32 (FcR) in BALB/c mouse footpad. Immunofluorescence staining for Nramp1 (green) and the macrophage-specific marker CD32 (FcR) (red) in the footpad cryosections of BALB/c mouse that were either uninfected or infected with wild type *L. major*. Tissues were harvested at 6 weeks post infection (p.i.). Nuclei were stained with DAPI (blue) and small *Leishmania* nuclei are marked with white arrows. The insets are zoomed regions marked by boxes showing co-localization of Nramp1 and CD32 (FcR). Images were acquired with Leica SP8 confocal, 63× objective.

**Table S1.** List of reagents used in the study.

| **Reagent** | **Source** | **Identifier** |
| --- | --- | --- |
| Adenine | Sigma-Aldrich | Cat# A2786 |
| Agarose | SRL | Cat#36601 |
| Ammonium acetate | Merck | Cat#101116 |
| Blasticidin S HCL | Gibco | Cat#R21001 |
| BSA | Sigma-Aldrich | Cat#A2153 |
| CaCl<sub>2</sub> | Sigma-Aldrich | Cat#C5080 |
| DMEM | Gibco | Cat#12100061 |
| EDTA | G-Biosciences | Cat#RC047 |
| FeCl<sub>3</sub> | Sigma-Aldrich | Cat#701122 |
| Ferrozine | Sigma-Aldrich | Cat#160601 |
| FBS | Gibco | Cat#97068-085 |
| Folic acid | Sigma-Aldrich | Cat#F8758 |
| Gentamicin | Abbott | N/A |
| Gelatin | AMRESCO | Cat#9764 |
| Glutaraldehyde | Sigma-Aldrich | Cat#G7776 |
| HEPES | Sigma-Aldrich | Cat#3784 |
| Hemin | Sigma-Aldrich | Cat# 51280 |
| HCL | Merck | Cat#1930010521 |
| KMnO<sub>4</sub> | Merck | Cat#105082 |
| L-Ascorbic acid | Sigma-Aldrich | Cat#A92902 |
| Lipofectamine 2000 tranfection reagent | Invitrogen | Cat#11668019 |
| M199 | Gibco | Cat#31100019 |
| NaCl | Merck | Cat# 106404 |
| NaHCO<sub>3</sub> | Sigma-Aldrich | Cat#S5761 |
| Neocuproine | Sigma-Aldrich | Cat#N1501 |
| Osmium tetroxide | Sigma-Aldrich | Cat#75632 |
| Optimal cutting temperature (OCT) compound | Tissue-Tek | Cat#4583 |
| Penicillin-streptomycin | Gibco | Cat#10378016 |
| Puromycin Dihydrochloride | Merck | Cat#P8833 |
| poly-L-lysine | Sigma-Aldrich | Cat#P4707 |
| polybrene | Sigma-Aldrich | Cat#TR-1003 |
| PMSF | Sigma-Aldrich | Cat# P7626 |
| PVDF | Millipore | Cat#IPVH00010 |
| Sodium pyruvate | Sigma-Aldrich | Cat#P5280 |
| SIGMAFAST Protease Inhibitor<br>Cocktail Tablet, EDTA-free | Sigma-Aldrich | Cat#S8830 |
| SDS | Sigma-Aldrich | Cat#L4390 |
| Super Signal West Pico<br>Chemiluminescent Substrate | Thermo Fisher Scientific | Cat#34580 |
| SYBR green fluorophore | BioRad | Cat#: 1725121 |
| Sucrose | Merck | Cat#1.94953.0521 |
| Skim Milk Powder | Millipore | Cat#70166 |
| Tween 20 | Sigma-Aldrich | Cat#93773 |
| Tris | Sigma-Aldrich | Cat#T6066 |
| TritonX -100 | Sigma-Aldrich | Cat#T9284 |
| Trypsin | Gibco | Cat#25200056 |
| TRIzol reagent | Invitrogen | Cat#15596026 |
| Vectashield | Vecta laboratories | Cat#H1000 |
| Zncl<sub>2</sub> | Merck | Cat#108816 |
| 1,10-Phen | Sigma-Aldrich | Cat#131377 |
| 0.22um Syringe filters | AXIVA | Cat#SFPS33R |
| Total Exosome Isolation Reagent | Invitrogen | Cat# 4478359 |
| Verso cDNA synthesis kit | Thermo Fisher Scientific | Cat# AB1453A |
| PureLink Quick Plasmid Miniprep Kit | Invitrogen | Cat# K210010 |
| DNase I | Thermo Fisher Scientific | Cat# 18068-015 |
| Phusion enzyme | Thermo Fisher Scientific | Cat# F530S |

**Table S2.**
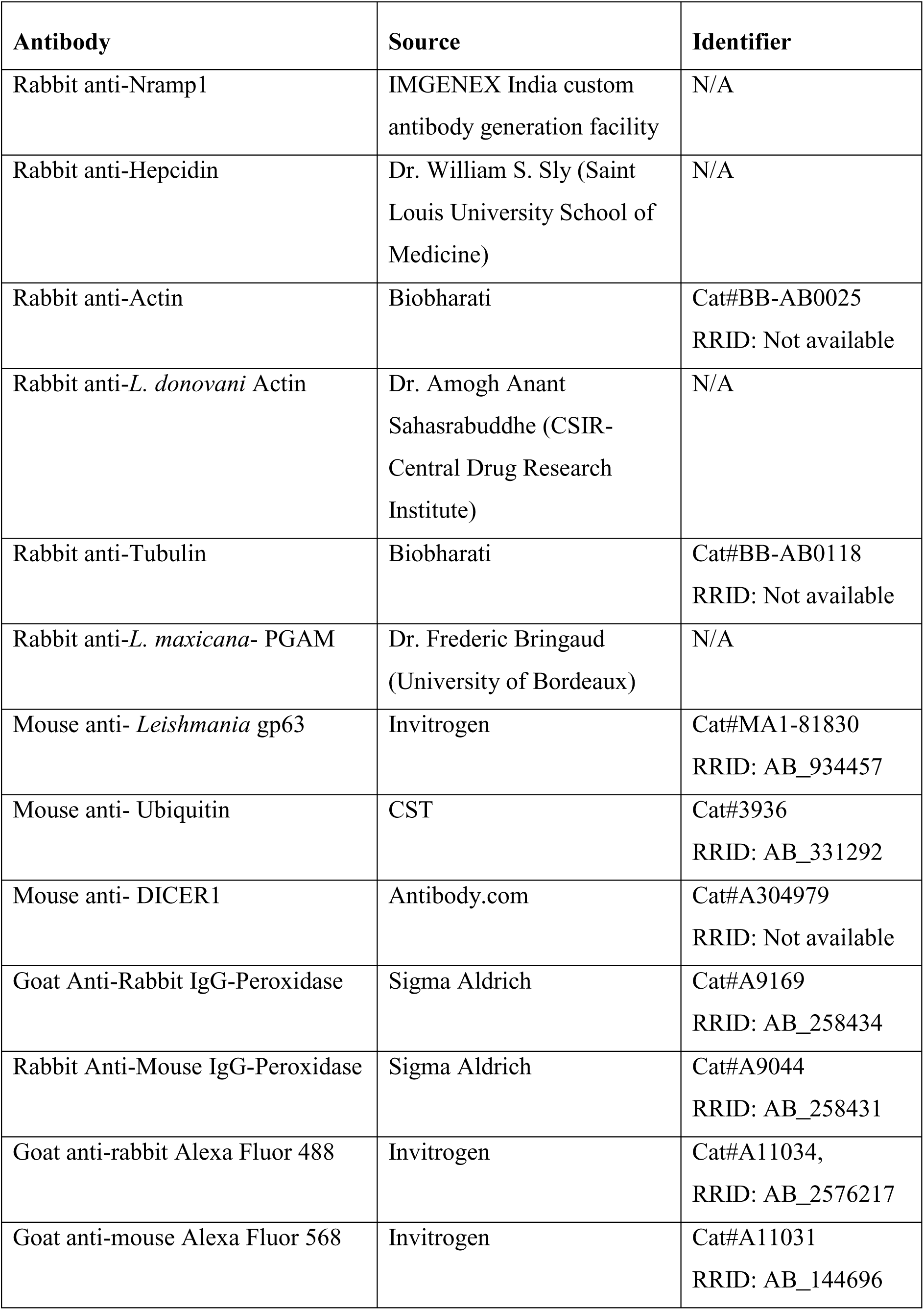
List of antibodies used in the study.

**Table S3.**
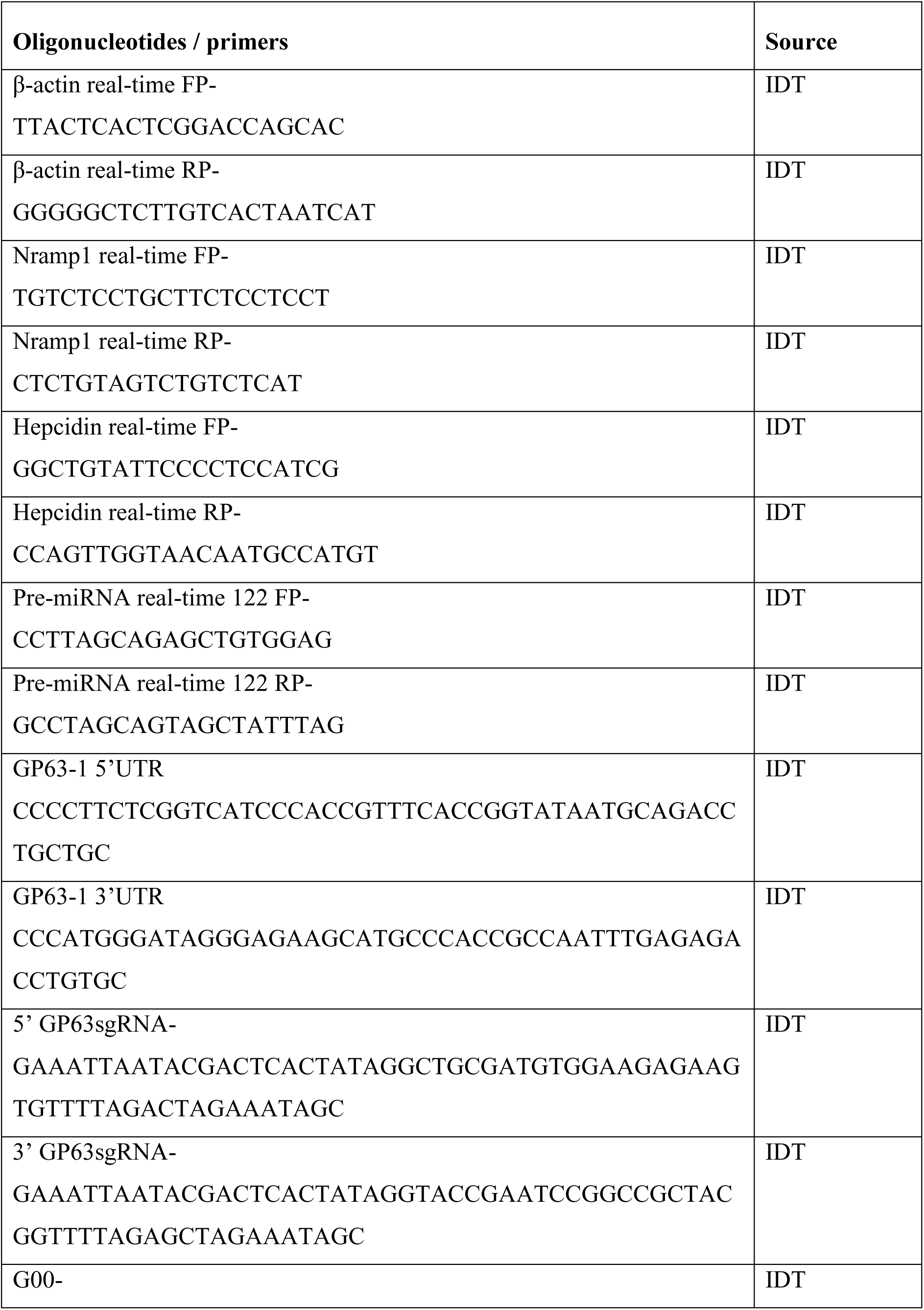
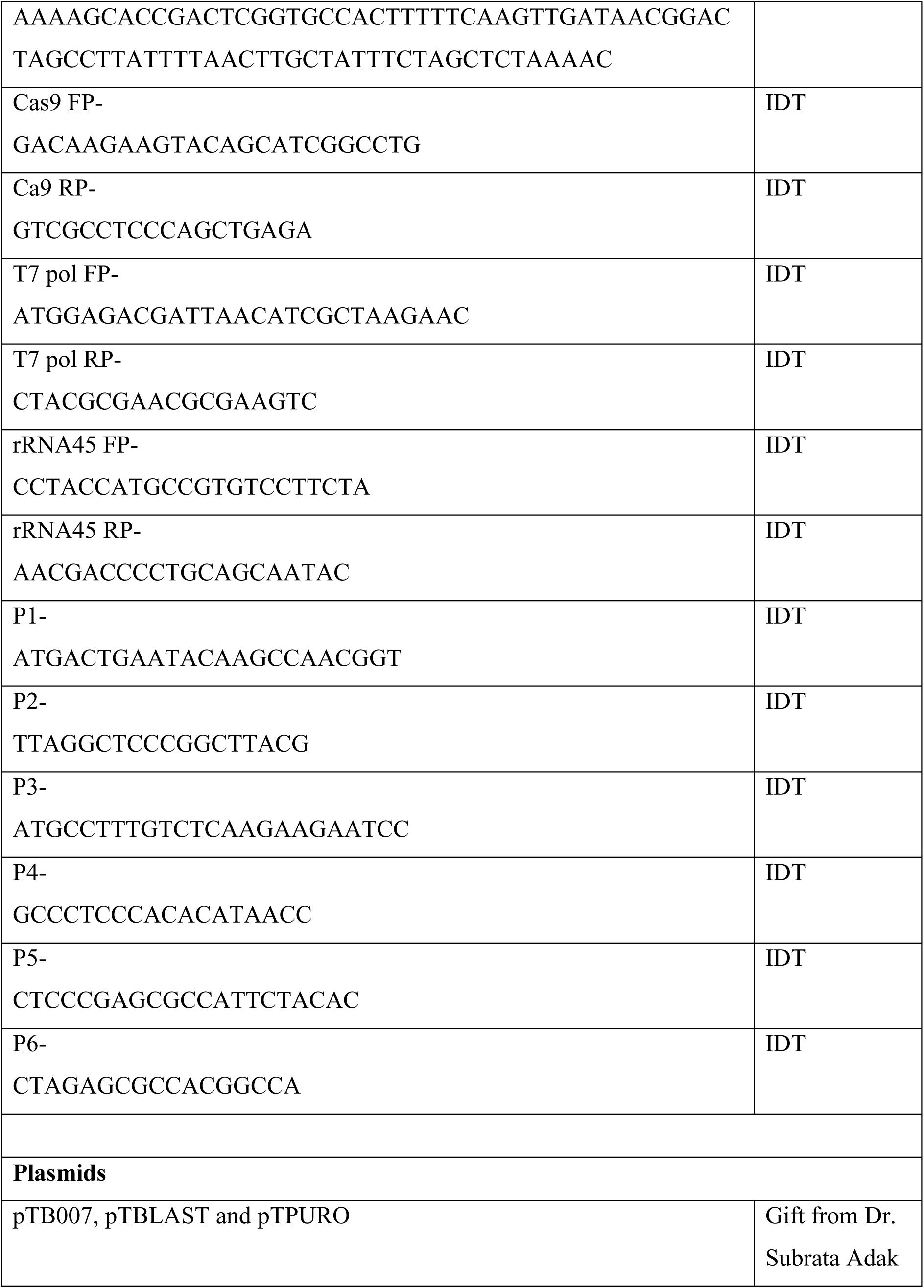
List of Oligonucleotides / primers and plasmids used in the study.

## REFERENCES

Alford, C.E., T.E. King Jr, and P.A. Campbell. 1991. Role of transferrin, transferrin receptors, and iron in macrophage listericidal activity. Journal of Experimental Medicine. 174:459–466. doi:10.1084/jem.174.2.459.

Alvar, J., I.D. Vélez, C. Bern, M. Herrero, P. Desjeux, J. Cano, J. Jannin, M. den Boer, and WHO Leishmaniasis Control Team. 2012. Leishmaniasis worldwide and global estimates of its incidence. PLoS One. 7:e35671. doi:10.1371/journal.pone.0035671.

Andriopoulos, B., E. Corradini, Y. Xia, S.A. Faasse, S. Chen, L. Grgurevic, M.D. Knutson, A. Pietrangelo, S. Vukicevic, H.Y. Lin, and J.L. Babitt. 2009. BMP6 is a key endogenous regulator of hepcidin expression and iron metabolism. Nat Genet. 41:482–487. doi:10.1038/ng.335.

Arango Duque, G., M. Fukuda, S.J. Turco, S. Stäger, and A. Descoteaux. 2014. Leishmania promastigotes induce cytokine secretion in macrophages through the degradation of synaptotagmin XI. J Immunol. 193:2363–2372. doi:10.4049/jimmunol.1303043.

Atkinson, P.G., and C.H. Barton. 1999. High level expression of Nramp1G169 in RAW264.7 cell transfectants: analysis of intracellular iron transport. Immunology. 96:656–662. doi:10.1046/j.1365-2567.1999.00672.x.

Banerjee, S., and R. Datta. 2020. Leishmania infection triggers hepcidin-mediated proteasomal degradation of Nramp1 to increase phagolysosomal iron availability. Cell Microbiol. 22:e13253. doi:10.1111/cmi.13253.

Banerjee, S., and R. Datta. 2023. Localized Leishmania major infection disrupts systemic iron homeostasis that can be controlled by oral iron supplementation. J Biol Chem. 299:105064. doi:10.1016/j.jbc.2023.105064.

Barton, C.H., T.E. Biggs, S.T. Baker, H. Bowen, and P.G.P. Atkinson. 1999. Nrampl: a link between intracellular iron transport and innate resistance to intracellular pathogens. Journal of Leukocyte Biology. 66:757–762. doi:10.1002/jlb.66.5.757.

Beneke, T., R. Madden, L. Makin, J. Valli, J. Sunter, and E. Gluenz. 2017. A CRISPR Cas9 high-throughput genome editing toolkit for kinetoplastids. R Soc Open Sci. 4:170095. doi:10.1098/rsos.170095.

Buracco, S., B. Peracino, R. Cinquetti, E. Signoretto, A. Vollero, F. Imperiali, M. Castagna, E. Bossi, and S. Bozzaro. 2015. Dictyostelium Nramp1, which is structurally and functionally similar to mammalian DMT1 transporter, mediates phagosomal iron efflux. J Cell Sci. 128:3304–3316. doi:10.1242/jcs.173153.

Burza, S., S.L. Croft, and M. Boelaert. 2018. Leishmaniasis. Lancet. 392:951–970. doi:10.1016/S0140-6736(18)31204-2.

Cairo, G., F. Bernuzzi, and S. Recalcati. 2006. A precious metal: Iron, an essential nutrient for all cells. Genes Nutr. 1:25–39. doi:10.1007/BF02829934.

Camaschella, C. 2009. BMP6 orchestrates iron metabolism. Nat Genet. 41:386–388. doi:10.1038/ng0409-386.

Cassat, J.E., and E.P. Skaar. 2013. Iron in infection and immunity. Cell Host Microbe. 13:509–519. doi:10.1016/j.chom.2013.04.010.

Castoldi, M., M. Vujic Spasic, S. Altamura, J. Elmén, M. Lindow, J. Kiss, J. Stolte, R. Sparla, L.A. D’Alessandro, U. Klingmüller, R.E. Fleming, T. Longerich, H.J. Gröne, V. Benes, S. Kauppinen, M.W. Hentze, and M.U. Muckenthaler. 2011. The liver-specific microRNA miR-122 controls systemic iron homeostasis in mice. J Clin Invest. 121:1386–1396. doi:10.1172/JCI44883.

Chang, K.-P., and D. Dwyer. 1978. Leishmania Donovani. Hamster macrophage interactions in vitro: cell entry, intracellular survival, and multiplication of amastigotes. J Exp Med. 147:515–530.

Chaudhuri, G., M. Chaudhuri, A. Pan, and K.P. Chang. 1989. Surface acid proteinase (gp63) of Leishmania mexicana. A metalloenzyme capable of protecting liposome-encapsulated proteins from phagolysosomal degradation by macrophages. J Biol Chem. 264:7483–7489.

Cheng, X., and H. Wang. 2012. Multiple targeting motifs direct NRAMP1 into lysosomes. Biochem Biophys Res Commun. 419:578–583. doi:10.1016/j.bbrc.2012.02.078.

Cunrath, O., and D. Bumann. 2019. Host resistance factor SLC11A1 restricts Salmonella growth through magnesium deprivation. Science. 366:995–999. doi:10.1126/science.aax7898.

Dixon, S.J., and B.R. Stockwell. 2014. The role of iron and reactive oxygen species in cell death. Nat Chem Biol. 10:9–17. doi:10.1038/nchembio.1416.

Drakesmith, H., E. Nemeth, and T. Ganz. 2015. Ironing out Ferroportin. Cell Metabolism. 22:777–787. doi:10.1016/j.cmet.2015.09.006.

Evans, C.A., M.S. Harbuz, T. Ostenfeld, A. Norrish, and J.M. Blackwell. 2001. Nramp1 is expressed in neurons and is associated with behavioural and immune responses to stress. Neurogenetics. 3:69–78. doi:10.1007/s100480100105.

Flannery, A.R., R.L. Renberg, and N.W. Andrews. 2013. Pathways of iron acquisition and utilization in Leishmania. Curr Opin Microbiol. 16:716–721. doi:10.1016/j.mib.2013.07.018.

Ghosh, J., M. Bose, S. Roy, and S.N. Bhattacharyya. 2013. Leishmania donovani targets Dicer1 to downregulate miR-122, lower serum cholesterol, and facilitate murine liver infection. Cell Host Microbe. 13:277–288. doi:10.1016/j.chom.2013.02.005.

Gomez, M.A., I. Contreras, M. Hallé, M.L. Tremblay, R.W. McMaster, and M. Olivier. 2009. Leishmania GP63 alters host signaling through cleavage-activated protein tyrosine phosphatases. Sci Signal. 2:ra58. doi:10.1126/scisignal.2000213.

Govoni, G., S. Gauthier, F. Billia, N.N. Iscove, and P. Gros. 1997. Cell-specific and inducible Nramp1 gene expression in mouse macrophages in vitro and in vivo. J Leukoc Biol. 62:277–286. doi:10.1002/jlb.62.2.277.

Guay-Vincent, M.-M., C. Matte, A.-M. Berthiaume, M. Olivier, M. Jaramillo, and A. Descoteaux. 2022. Revisiting Leishmania GP63 host cell targets reveals a limited spectrum of substrates. PLoS Pathog. 18:e1010640. doi:10.1371/journal.ppat.1010640.

Haas, A. 2007. The phagosome: compartment with a license to kill. Traffic. 8:311–330. doi:10.1111/j.1600-0854.2006.00531.x.

Harrison, P.M., and P. Arosio. 1996. The ferritins: molecular properties, iron storage function and cellular regulation. Biochim Biophys Acta. 1275:161–203. doi:10.1016/0005-2728(96)00022-9.

Hassani, K., E. Antoniak, A. Jardim, and M. Olivier. 2011. Temperature-induced protein secretion by Leishmania mexicana modulates macrophage signalling and function. PLoS One. 6:e18724. doi:10.1371/journal.pone.0018724.

Hassani, K., and M. Olivier. 2013. Immunomodulatory impact of leishmania-induced macrophage exosomes: a comparative proteomic and functional analysis. PLoS Negl Trop Dis. 7:e2185. doi:10.1371/journal.pntd.0002185.

Hassani, K., M.T. Shio, C. Martel, D. Faubert, and M. Olivier. 2014. Absence of metalloprotease GP63 alters the protein content of Leishmania exosomes. PLoS One. 9:e95007. doi:10.1371/journal.pone.0095007.

Hefnawy, A., M. Berg, J.-C. Dujardin, and G. De Muylder. 2017. Exploiting Knowledge on *Leishmania* Drug Resistance to Support the Quest for New Drugs. Trends in Parasitology. 33:162–174. doi:10.1016/j.pt.2016.11.003.

Huynh, C., D.L. Sacks, and N.W. Andrews. 2006. A Leishmania amazonensis ZIP family iron transporter is essential for parasite replication within macrophage phagolysosomes. J Exp Med. 203:2363–2375. doi:10.1084/jem.20060559.

Isnard, A., M.T. Shio, and M. Olivier. 2012. Impact of Leishmania metalloprotease GP63 on macrophage signaling. Front Cell Infect Microbiol. 2:72. doi:10.3389/fcimb.2012.00072.

Ivens, A.C., C.S. Peacock, E.A. Worthey, L. Murphy, G. Aggarwal, M. Berriman, E. Sisk, M.-A. Rajandream, E. Adlem, R. Aert, A. Anupama, Z. Apostolou, P. Attipoe, N. Bason, C. Bauser, A. Beck, S.M. Beverley, G. Bianchettin, K. Borzym, G. Bothe, C.V. Bruschi, M. Collins, E. Cadag, L. Ciarloni, C. Clayton, R.M.R. Coulson, A. Cronin, A.K. Cruz, R.M. Davies, J. De Gaudenzi, D.E. Dobson, A. Duesterhoeft, G. Fazelina, N. Fosker, A.C. Frasch, A. Fraser, M. Fuchs, C. Gabel, A. Goble, A. Goffeau, D. Harris, C. Hertz-Fowler, H. Hilbert, D. Horn, Y. Huang, S. Klages, A. Knights, M. Kube, N. Larke, L. Litvin, A. Lord, T. Louie, M. Marra, D. Masuy, K. Matthews, S. Michaeli, J.C. Mottram, S. MuÌ^ller-Auer, H. Munden, S. Nelson, H. Norbertczak, K. Oliver, S. O’Neil, M. Pentony, T.M. Pohl, C. Price, B. Purnelle, M.A. Quail, E. Rabbinowitsch, R. Reinhardt, M. Rieger, J. Rinta, J. Robben, L. Robertson, J.C. Ruiz, S. Rutter, D. Saunders, M. SchaÌ^fer, J. Schein, D.C. Schwartz, K. Seeger, A. Seyler, S. Sharp, H. Shin, D. Sivam, R. Squares, S. Squares, V. Tosato, C. Vogt, G. Volckaert, R. Wambutt, T. Warren, H. Wedler, J. Woodward, S. Zhou, W. Zimmermann, D.F. Smith, J.M. Blackwell, et al. 2005. The Genome of the Kinetoplastid Parasite, Leishmania major. Science. 309:436–442. doi:10.1126/science.1112680.

Jabado, N., A. Jankowski, S. Dougaparsad, V. Picard, S. Grinstein, and P. Gros. 2000. Natural resistance to intracellular infections: natural resistance-associated macrophage protein 1 (Nramp1) functions as a pH-dependent manganese transporter at the phagosomal membrane. J Exp Med. 192:1237–1248. doi:10.1084/jem.192.9.1237.

Joshi, P.B., B.L. Kelly, S. Kamhawi, D.L. Sacks, and W.R. McMaster. 2002. Targeted gene deletion in Leishmania major identifies leishmanolysin (GP63) as a virulence factor. Mol Biochem Parasitol. 120:33–40. doi:10.1016/s0166-6851(01)00432-7.

Livak, K.J., and T.D. Schmittgen. 2001. Analysis of relative gene expression data using real-time quantitative PCR and the 2(-Delta Delta C(T)) Method. Methods. 25:402–408. doi:10.1006/meth.2001.1262.

Marquis, J.-F., and P. Gros. 2007. Intracellular Leishmania: your iron or mine? Trends Microbiol. 15:93–95. doi:10.1016/j.tim.2007.01.001.

Matheoud, D., N. Moradin, A. Bellemare-Pelletier, M.T. Shio, W.J. Hong, M. Olivier, E. Gagnon, M. Desjardins, and A. Descoteaux. 2013. Leishmania evades host immunity by inhibiting antigen cross-presentation through direct cleavage of the SNARE VAMP8. Cell Host Microbe. 14:15–25. doi:10.1016/j.chom.2013.06.003.

McConville, M.J., D. de Souza, E. Saunders, V.A. Likic, and T. Naderer. 2007. Living in a phagolysosome; metabolism of Leishmania amastigotes. Trends Parasitol. 23:368–375. doi:10.1016/j.pt.2007.06.009.

McGwire, B.S., W.A. O’Connell, K.-P. Chang, and D.M. Engman. 2002. Extracellular release of the glycosylphosphatidylinositol (GPI)-linked Leishmania surface metalloprotease, gp63, is independent of GPI phospholipolysis: implications for parasite virulence. J Biol Chem. 277:8802–8809. doi:10.1074/jbc.M109072200.

Muckenthaler, M.U., S. Rivella, M.W. Hentze, and B. Galy. 2017. A Red Carpet for Iron Metabolism. Cell. 168:344–361. doi:10.1016/j.cell.2016.12.034.

Murdoch, C.C., and E.P. Skaar. 2022. Nutritional immunity: the battle for nutrient metals at the host–pathogen interface. Nat Rev Microbiol. 20:657–670. doi:10.1038/s41579-022-00745-6.

Nairz, M., A. Schroll, T. Sonnweber, and G. Weiss. 2010. The struggle for iron - a metal at the host-pathogen interface. Cell Microbiol. 12:1691–1702. doi:10.1111/j.1462-5822.2010.01529.x.

Nemeth, E., M.S. Tuttle, J. Powelson, M.B. Vaughn, A. Donovan, D.M. Ward, T. Ganz, and J. Kaplan. 2004. Hepcidin Regulates Cellular Iron Efflux by Binding to Ferroportin and Inducing Its Internalization. Science. 306:2090–2093. doi:10.1126/science.1104742.

Pal, D.S., M. Abbasi, D.K. Mondal, B.A. Varghese, R. Paul, S. Singh, and R. Datta. 2017. Interplay between a cytosolic and a cell surface carbonic anhydrase in pH homeostasis and acid tolerance of Leishmania. J Cell Sci. 130:754–766. doi:10.1242/jcs.199422.

Peters, N.C., J.G. Egen, N. Secundino, A. Debrabant, N. Kimblin, S. Kamhawi, P. Lawyer, M.P. Fay, R.N. Germain, and D. Sacks. 2008. In vivo imaging reveals an essential role for neutrophils in leishmaniasis transmitted by sand flies. Science. 321:970–974. doi:10.1126/science.1159194.

Pissarra, J., J. Pagniez, E. Petitdidier, M. Séveno, O. Vigy, R. Bras-Gonçalves, J.-L. Lemesre, and P. Holzmuller. 2022. Proteomic Analysis of the Promastigote Secretome of Seven Leishmania Species. J Proteome Res. 21:30–48. doi:10.1021/acs.jproteome.1c00244.

Ponte-Sucre, A., F. Gamarro, J.-C. Dujardin, M.P. Barrett, R. López-Vélez, R. García-Hernández, A.W. Pountain, R. Mwenechanya, and B. Papadopoulou. 2017. Drug resistance and treatment failure in leishmaniasis: A 21st century challenge. PLoS Negl Trop Dis. 11:e0006052. doi:10.1371/journal.pntd.0006052.

Russell, D.G., and H. Wilhelm. 1986. The involvement of the major surface glycoprotein (gp63) of Leishmania promastigotes in attachment to macrophages. J Immunol. 136:2613–2620.

Santarém, N., G. Racine, R. Silvestre, A. Cordeiro-da-Silva, and M. Ouellette. 2013. Exoproteome dynamics in Leishmania infantum. J Proteomics. 84:106–118. doi:10.1016/j.jprot.2013.03.012.

Searle, S., N.A. Bright, T.I. Roach, P.G. Atkinson, C.H. Barton, R.H. Meloen, and J.M. Blackwell. 1998. Localisation of Nramp1 in macrophages: modulation with activation and infection. J Cell Sci. 111 ( Pt 19):2855–2866. doi:10.1242/jcs.111.19.2855.

Sen, S., S.K. Bal, S. Yadav, P. Mishra, V.V. G, R. Rastogi, and C.K. Mukhopadhyay. 2022. Intracellular pathogen Leishmania intervenes in iron loading into ferritin by cleaving chaperones in host macrophages as an iron acquisition strategy. J Biol Chem. 298:102646. doi:10.1016/j.jbc.2022.102646.

Seth, A., A. Das, and R. Datta. 2025. Identification of a basal body-localized epsilon-tubulin in Leishmania. FEBS Lett. doi:10.1002/1873-3468.70012.

Silverman, J.M., J. Clos, C.C. de’Oliveira, O. Shirvani, Y. Fang, C. Wang, L.J. Foster, and N.E. Reiner. 2010. An exosome-based secretion pathway is responsible for protein export from Leishmania and communication with macrophages. J Cell Sci. 123:842–852. doi:10.1242/jcs.056465.

Soe-Lin, S., S.S. Apte, B. Andriopoulos, M.C. Andrews, M. Schranzhofer, T. Kahawita, D. Garcia-Santos, and P. Ponka. 2009. Nramp1 promotes efficient macrophage recycling of iron following erythrophagocytosis in vivo. Proceedings of the National Academy of Sciences. 106:5960–5965. doi:10.1073/pnas.0900808106.

Soe-Lin, S., A.D. Sheftel, B. Wasyluk, and P. Ponka. 2008. Nramp1 equips macrophages for efficient iron recycling. Exp Hematol. 36:929–937. doi:10.1016/j.exphem.2008.02.013.

Taylor, M.C., and J.M. Kelly. 2010. Iron metabolism in trypanosomatids, and its crucial role in infection. Parasitology. 137:899–917. doi:10.1017/S0031182009991880.

Vidal, S., P. Gros, and E. Skamene. 1995. Natural resistance to infection with intracellular parasites: molecular genetics identifies Nramp1 as the Bcg/Ity/Lsh locus. J Leukoc Biol. 58:382–390. doi:10.1002/jlb.58.4.382.

Vidal, S.M., D. Malo, K. Vogan, E. Skamene, and P. Gros. 1993. Natural resistance to infection with intracellular parasites: isolation of a candidate for Bcg. Cell. 73:469–485. doi:10.1016/0092-8674(93)90135-d.

Vidal, S.M., E. Pinner, P. Lepage, S. Gauthier, and P. Gros. 1996. Natural resistance to intracellular infections: Nramp1 encodes a membrane phosphoglycoprotein absent in macrophages from susceptible (Nramp1 D169) mouse strains. The Journal of Immunology. 157:3559–3568. doi:10.4049/jimmunol.157.8.3559.

Wang, R.-H., C. Li, X. Xu, Y. Zheng, C. Xiao, P. Zerfas, S. Cooperman, M. Eckhaus, T. Rouault, L. Mishra, and C.-X. Deng. 2005. A role of SMAD4 in iron metabolism through the positive regulation of hepcidin expression. Cell Metab. 2:399–409. doi:10.1016/j.cmet.2005.10.010.

Wu, X.-G., Y. Wang, Q. Wu, W.-H. Cheng, W. Liu, Y. Zhao, C. Mayeur, P.J. Schmidt, P.B. Yu, F. Wang, and Y. Xia. 2014. HFE interacts with the BMP type I receptor ALK3 to regulate hepcidin expression. Blood. 124:1335–1343. doi:10.1182/blood-2014-01-552281.

Zhang, X., R. Goncalves, and D.M. Mosser. 2008. The isolation and characterization of murine macrophages. Curr Protoc Immunol. Chapter 14:14.1.1-14.1.14. doi:10.1002/0471142735.im1401s83.

